# The Backbone Network of Dynamic Functional Connectivity

**DOI:** 10.1101/2021.04.20.440711

**Authors:** Nima Asadi, Ingrid R. Olson, Zoran Obradovic

## Abstract

Temporal networks have become increasingly pervasive in many real-world applications, including the functional connectivity analysis of spatially separated regions of the brain. A major challenge in analysis of such networks is the identification of noise confounds, which introduce temporal ties that are non-essential, or links that are formed by chance due to local properties of the nodes. Several approaches have been suggested in the past for static networks or temporal networks with binary weights for extracting significant ties whose likelihood cannot be reduced to the local properties of the nodes. In this work, we propose a data-driven procedure to reveal the irreducible ties in dynamic functional connectivity of resting state fRMI data with continuous weights. This framework includes a null model that estimates the latent characteristics of the distributions of temporal links through optimization, followed by a statistical test to filter the links whose formation can be reduced to the activities and local properties of their interacting nodes. We demonstrate the benefits of this approach by applying it to a resting state fMRI dataset, and provide further discussion on various aspects and advantages of it.

## 2. Introduction

Dynamic functional connectivity (dFC) has been widely used to analyze temporal associations among separate regions of the brain as well as the correlation between functional patterns of connectivity and cognitive abilities (Malsburg, Phillps, and Singer 2010; Allen et al. 2014; Van Dijk, Hedden, et al. 2010; Jones et al. 2012). In order to identify co-activation patterns of dFC over the period of experiment, a temporal segmentation (such as sliding window) is commonly applied on the time courses of BOLD activation of brain regions to divide them into consecutive temporal windows (Smith et al. 2012; Hutchison et al. 2013; Allen et al. 2014). Then, the connectivity between separate regions is measured to generate one graph adjacency matrix per each temporal window (Damaraju et al. 2014). Building on this core framework, several enhancements have been proposed in the past years, such as different temporal segmentation approaches, to increase the power and precision of dFC analysis (Chang and Glover 2010; Kiviniemi et al. 2011; Heitmann and Breakspear 2018; Hindriks et al. 2016).

However, a major challenge in analysis of dynamic functional connectivity is to distinguish and address the existing noise confounds in the data, which in turn influences the brain connectivity measures and the structure of the dFC network (Birn et al. 2008; Shmueli et al. 2007; Chang and Glover 2009; Kalthoff et al. 2011). This issue especially intensifies with the increase in spatial resolution of the analysis, i.e. in smaller regions of interest or voxel-level connectivity, as well as in resting state fMRI data (Birn 2012; Kalthoff et al. 2011; Hallquist, Hwang, and Luna 2013). There are several possible sources of noise in resting state fMRI data, including displacements, even as small as a millimeter or less, which could add random noise to the generated time series, and therefore decrease the statistical power in resting state functional connectivity (rsFC) analysis (Van Dijk, Sabuncu, and Buckner 2012). Even more challenging, it can result in false positive or negative activation if the displacements are correlated with the stimuli (Lydon-Staley et al. 2019; Patanaik et al. 2018; Savva et al. 2020). Cardiovascular and respiratory signals are also widely identified as a source of noise, causing synchronized fluctuations in MRI signal (Glover and Lee 1995). This type of noise could vary based on individual differences and potentially also become temporally synchronized to the stimulus. Hemodynamic lags could cause mis-localisation of cortical activation and statistical methods needed optimisation for the formidable problem of multiple comparisons in typical fMRI datasets (Glover and Lee 1995; Tong, Hocke, and Frederick 2019).

Due to these challenges, neuroscientists often face the concern of analytical models being noise-induced (K. Murphy, Birn, and Bandettini 2013; Gorgolewski et al. 2013; Choe et al. 2017). A number of correction techniques have been suggested in the past to reduce the influence of these confounds, including modelling fMRI signal variations using independent measures of the cardiac and respiratory signal variations (Bright and K. Murphy 2017; K. Murphy, Birn, and Bandettini 2013; Bollmann et al. 2017; Behzadi et al. 2007). However, the effect of various sources of noise on the dynamic connectivity of fMRI data is yet to be addressed through a data-driven and systematic framework (Kundu et al. 2012; Beall and Lowe 2007).

Moreover, temporal ties that can be reduced to node properties can exist between nodes due to the nature of the data itself. Highly active regions could in principle form a larger number of trivial ties with other regions, and reciprocally, the information of ties that regions with lower activity form can be lost in common analytical procedures (Kobayashi, Takaguchi, and Barrat 2019; Gemmetto, Cardillo, and Garlaschelli 2017). In general, if the network representation of a real-world system can be inferred based on local properties of the nodes, such as their activity level or degree, the true interaction and functional homologies between the nodes can not be detected (Gemmetto, Cardillo, and Garlaschelli 2017). It is therefore essential to identify truly dyadic relationships in the network in order to identify the non-redundant representation of the system, which in this study is the resting state functional connectivity of brain regions.

Therefore, the objective of this work is to put forward a data-driven approach to distinguish the significant ties that construct the functional connectivity of the brain from ties that are the result of random observational errors or chance. The latter group of temporal links are known as reducible ties, whereby they can be fully attributed to intrinsic node-specific features such as degree or strength of their link weights. On the other hand, the temporal ties that are incompatible with the null hypothesis of links being produced at random are known as irreducible or significant ties, and the network of such significant ties is known as the backbone network. Therefore, the goal of this study is to develop a data-driven framework to infer the two-dimensional backbone network from the multilayer network of dynamic functional connectivity.

Multiple approaches have been proposed to extract the significant ties in a network through statistical means, most of which target static networks (Alvarez-Hamelin et al. 2005; Ma et al. 2016; Serrano, Boguná, and Vespignani 2009; Tumminello et al. 2011; Gemmetto, Cardillo, and Garlaschelli 2017; Casiraghi et al. 2017; Kobayashi, Takaguchi, and Barrat 2019; X. Yan et al. 2018; Nadini et al. 2020). Across these approaches, a key step towards inferring the backbone network is the formulation of a reliable null model to characterize the reducible fraction of the temporal interactions, and to steer the procedure of filtering that fraction of network links. Several null models have been suggested in the literature whose focus is on static networks with continuous and binary weights, from basic weight thresholding of multilayer networks to more advanced techniques (Tumminello et al. 2011; M.-X. Li et al. 2014; Cimini et al. 2019; Kobayashi, Takaguchi, and Barrat 2019).

One of the main disadvantages with weight thresholding approaches is that they commonly fail to control for the difference in intrinsic attributes of the nodes, thus favor highly active nodes or nodes with other strong local properties, which can potentially have a large number of reducible links. A backbone approach for dynamic networks was proposed by Kobayashi et. al, in which they calculate the latent variables called activities that drive the probability of forming connection between nodes through the maximum likelihood estimation (MLE) approach (Kobayashi, Takaguchi, and Barrat 2019). That work is based on a network modeling concept named activity-driven networks (ADNs) where individual propensity of generating connections over time is determined by a latent nodal parameter commonly known as activity, and the probability of creating a link at a specific time instant between two nodes is the product of the individual latent activities of interacting nodes (Perra et al. 2012; Zino, Rizzo, and Porfiri 2017; Starnini and Pastor-Satorras 2014). Due to their analytical flexibility and interpretability, activity-driven network models have gained popularity in explaining features of real networks in various areas of research (Zino, Rizzo, and Porfiri 2017; Liu et al. 2014; Rizzo, Frasca, and Porfiri 2014; Zino, Rizzo, and Porfiri 2017). However, in mentioned studies, a binomial or Poisson distribution is considered for the temporal connections over time, which limits the approach to unweighted networks (Kobayashi, Takaguchi, and Barrat 2019; Nadini et al. 2020). Many relational networks based on real data, including various types of fMRI-based networks have continuous weights which contain significant information regarding the interactions between the nodes as well as the local and global properties of the network. These weights are commonly calculated based on statistical methods such as the correlation between the BOLD activation levels of the brainregions of interest or voxels. Therefore, inspired by the work of Kobayashi et. al, we propose an approach for extracting the significant ties for networks with continuous weights, which is appropriate for dynamic functional connectivity among many other applications with dynamic weighted networks. Obviously, this approach can be used for other weighted temporal networks which meet the characteristics of normality and independence of temporal ties, which are discussed in the methodology section. We demonstrate that this methodology controls for intrinsic local node attributes, with a null model that not only takes into account the global structure of the network, but also the temporal variations of the dynamic connectivity links. In the next section we explain the proposed approach in detail, followed by the experimental results on a real dataset of resting state fMRI. We then present an analysis of the results and discuss the advantages and shortcomings of our approach.

## 3. Methodology

In this section, we outline the methodological framework for identifying the significant links from the networks of dynamic resting state functional connectivity. A key step towards extracting the backbone network is the formulation of a valid and robust null model. For the sake of simplicity, we name our proposed approach the weighted backbone network (WBN). A null model assumes that all connections are formed randomly, meaning that the probability of a interaction between two nodes at a specific time window and the weight of interactions between them could be explained by chance (Kobayashi, Takaguchi, and Barrat 2019; Nadini et al. 2020; Gemmetto, Cardillo, and Garlaschelli 2017). The objective of inferring the backbone network is thus to detect links that are not compatible with the null hypothesis, meaning that their occurrence or strength is not driven by chance.

The null model that we present can be interpreted as a temporal fitness model, which is characterized by latent parameters that shape its distribution. In this vein, the first step is to estimate these parameters which are not directly observed from the data. For this purpose, we use a maximum likelihood estimation approach that exploits the global and temporal information of the network of dynamic connectivity. We discuss the details of this methodology in the next section.

### 3.1. Estimation of latent distribution variables

We consider a dynamic network of *N* nodes with links evolving over *τ* observation windows of size Δ such that *τ* =1,…, *τ*. At each time step *t*, a weighted undirected network is formed whose adjacency matrix *A_t_* stochastically varies in time, and the weights of temporal links (links that are formed at time step *t*) between each pair of nodes *i* and *j* form a Gaussian distribution over the *τ* time steps. Normality of the distribution of weights between each pair of nodes over time *τ* is concluded based on Central Limit Theorem and the assumption that the distribution of temporal weights has a finite variance (Dudley 1978; Dudley 2014; Haller and Bartsch 2009; Smith 2012). Moreover, an empirical assessment of normality of the distribution of temporal weights on a real dataset of resting state fMRI is provided in the result section.

We define a temporal null model in which each node *i* is assigned two intrinsic variables *ai,bi* ∈ (0,1], that rule the probability of mean *μ* and standard deviation *σ* of the temporal distribution of its interactions with other nodes over *τ* time steps, such that:

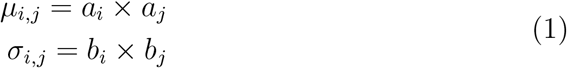

Therefore, a each parameter of the distributions of temporal ties between each pair of node *i* and *j* is the realization of a Bernoulli variable. The null model thus lays out a baseline for the expected mean and standard deviation of the distribution of interactions between two nodes over *τ* time given their intrinsic variables, if interacting nodes are selected at random at each time step.

To uncover significant links with regards to the null model described above, we proceed in two steps. First, given a set of weighted undirected temporal networks with *N* nodes, we estimate the intrinsic variables 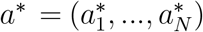 and 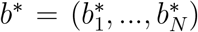 by calculating the maximum likelihood estimation of the set of parameters for each node. For this purpose, we consider the joint probability function over *τ* time intervals and edge weights 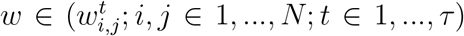 for the entire temporal connections of the network. Therefore we have:

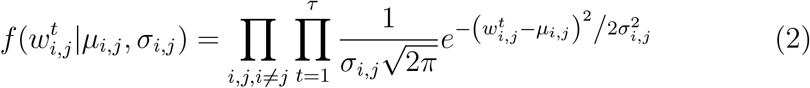

Where *μ_i,j_* and *σ_i,j_* denote the mean and standard deviation of the distribution of temporal edges between nodes *i* and *j* observed over *τ* time steps in the null model.

The log-likelihood function for the empirical data 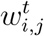 (weight of the link between *i* and *j* at time interval *t*) with replacing the values of *μ_i,j_* = *a_i_.a_j_* and *σ_i,j_* = *b_i_.b_j_* will lead to:

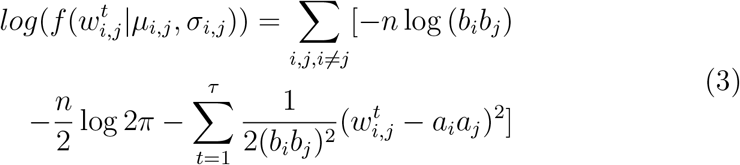

By differentiating the log-likelihood function with respect to the first parameter, *a_i_*, and setting it to zero we have:

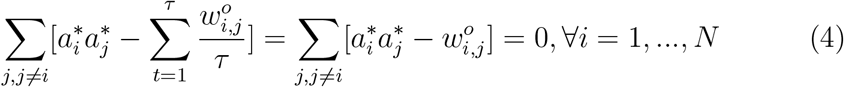

Similarly, by differentiating the log-likelihood function with respect to *b_i_* and setting it to zero we have:

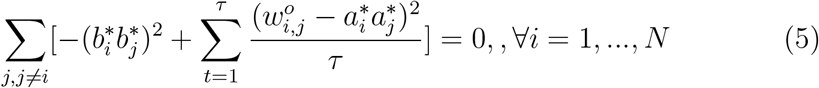

Where the the maximum likelihood estimation of 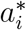 for every node *i* which was calculated from equation 4. Therefore, for a temporal network with *N* nodes, the pair of latent variables *a_i_, b_i_* for each node *i* can be estimated by solving the system of *N* nonlinear equations 5 and 6. The system of nonlinear equations can be solved through a standard numerical algorithm such as the newton method. The initial values of *a_i_* and *b_i_* are calculated by dividing the temporal degree of node *i* averaged over *τ* time steps by the doubled number of total temporal edges as follows:

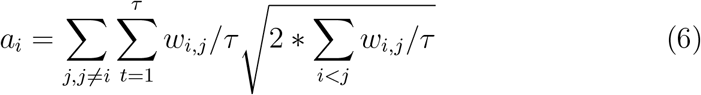

The general schema of the proposed methodology is provided in figure 1. Note that the proposed maximum likelihood approach incorporates the global information of the network as the weights of all temporal links are considered in the system of equations, and the estimation of latent activity levels is not merely dependent on local characteristics such as nodal degree or centrality measures.

**Figure 1:**
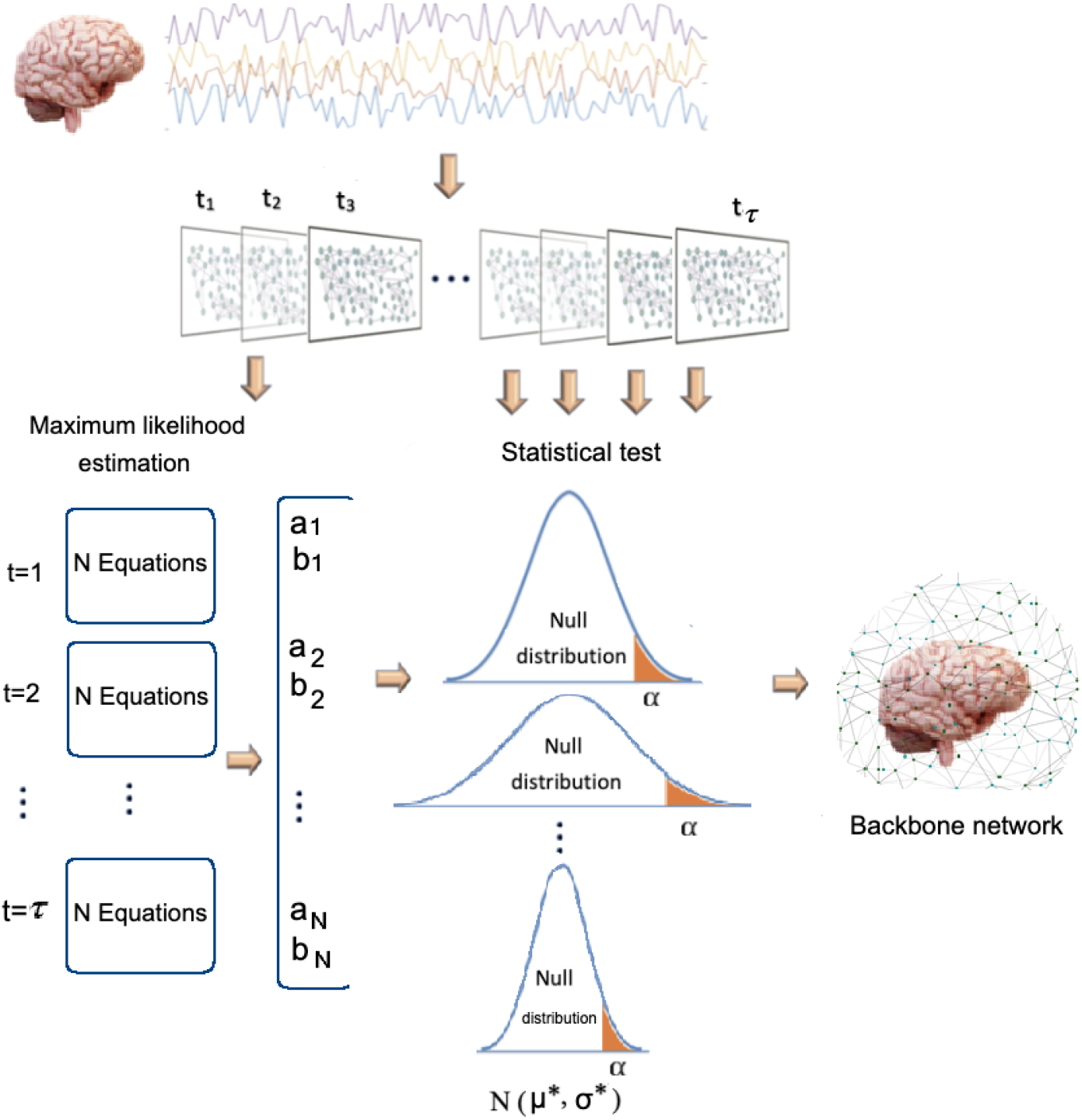
A schema of the Backbone network inference procedure

### 3.2. Selection of significant ties

After estimating the latent distribution variables for each node, we then compute for each pair of nodes *i* and *j* the probability distribution of their interaction over the *τ* time steps in the null model:

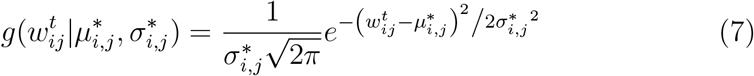

where 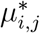 and 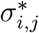 are the mean and standard deviation based on the estimated latent variables *a_i_, b_i_, a_j_, b_j_* through maximum likelihood estimation and equation 1. In order to determine the reducibility of a temporal link 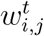 between nodes *i* and *j* at time *t*, it is compared against the *c*-th percentile (0 ≤ *c* ≤ 100) weight 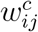 of the maximum likelihood estimated distribution of temporal links between *i* and *j*. If the empirical value 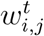 is larger than the 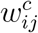, then it cannot be explained by the null model at significance level 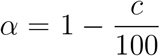. Therefore, the link 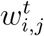 is determined to be a significant tie. The significance level *α* is given as an input to the model, providing a systematic adjustable filtering mechanism. The significance threshold *α* can also be assigned with Bonferroni correction, in which it can be adjusted by dividing by the sum of weights of edges to control for false positives. The P-value of the test is thus given by:

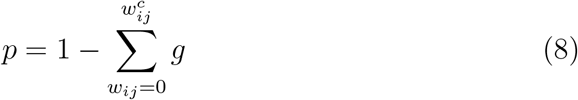

The reducibility of any temporal link within any time window *t* can be examined through this approach.

In order to determine whether a significant tie *w_ij_* exists between nodes *i* and *j* in the final backbone network, we simply count the number of times that the temporal link between them was found to be significant according to the threshold value *α*. If the count of significant links between *i* and *j* falls above half of the *τ* time intervals, i.e. 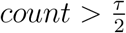, the link *w_ij_* is retained in the backbone network. Note that the final backbone network is a binary network, meaning that the weight of links is 1 if the link between two nodes is determined to be significant, and 0 otherwise. However, a weighted network of significant ties can be easily created through various error measures such as averaging the difference between the weights of the temporal links 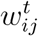 and the *c*-th percentile weight 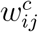 of the distribution.

An important property of the proposed null model is that the tie between two nodes at time *t* can be significant even if the weight of temporal link 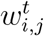 is small, with the condition that their individual latent variables *a, b*, and in turn the mean and standard deviation of their temporal distribution, are sufficiently low. On the contrary, ties with large weights might not be deemed to be significant by WBN if the estimated *a, b* corresponding to the two involved nodes are large. This property is illustrated in figure 2, where large estimated *μ* = *a_i_.a_j_* shifts the *c* – *th* percentile threshold to the right side of the distribution such that it becomes increasingly difficult for temporal links to meet the threshold. On the other hand, a small *a_i_.a^j^* makes it possible for links with smaller weights to be admitted into the backbone network due to the shift in *c* – *th* percentile threshold to the left.

**Figure 2:**
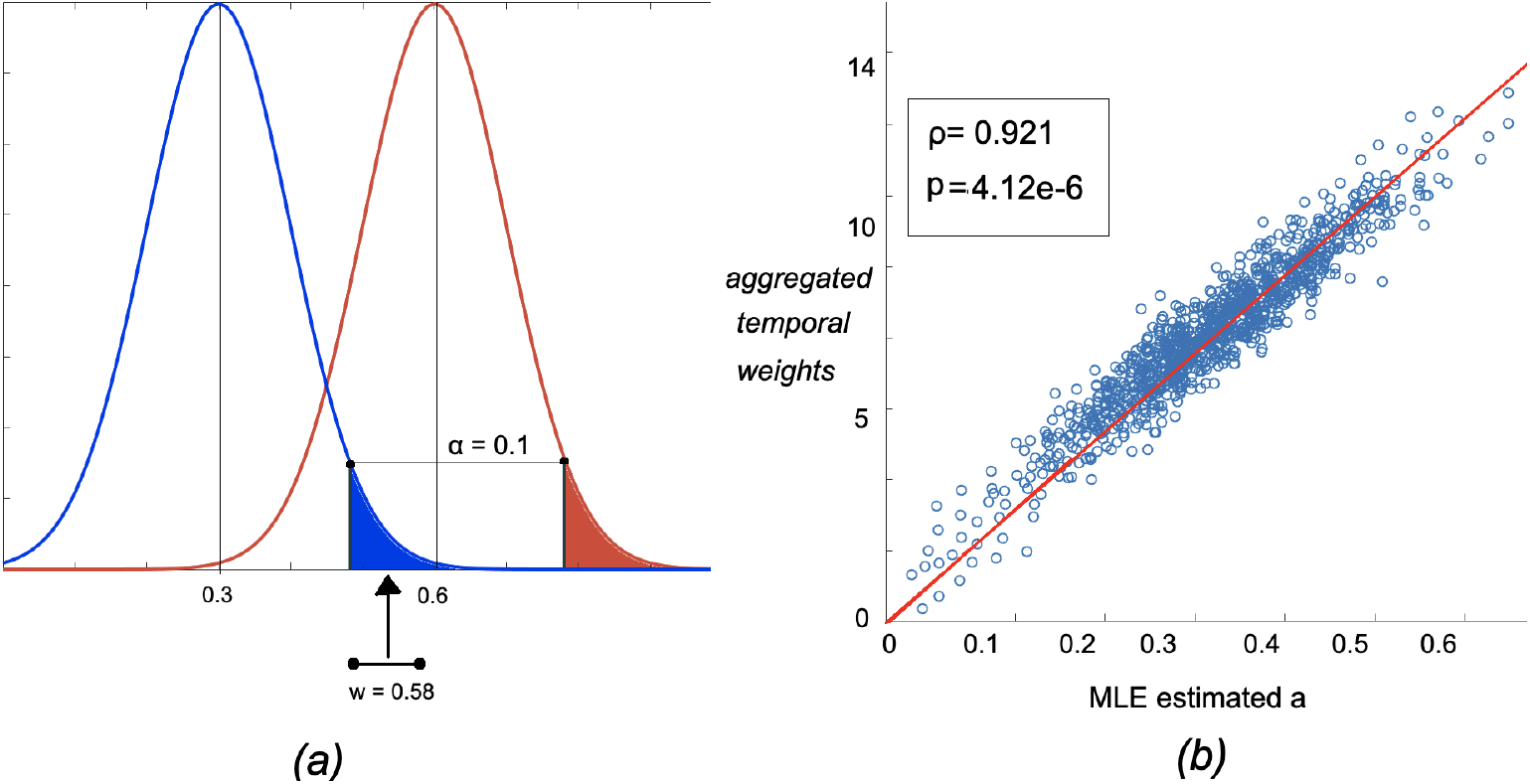
Figure a: an example of the effect of estimated *a* (distribution mean parameter), on admissibility of an empirical temporal link *w* where the threshold *α* is 0.1 (90th percentile). A link with a low weight can be admitted to the backbone network as long as its estimated mean is sufficiently low (the blue distribution), therefore, controlling for the effect of high intrinsic distribution weights on acceptance to the backbone network. Figure b: correlation between estimated latent distribution mean variable for each node *i* (*a_i_*) and the aggregated dFC weights corresponding to the node over *τ* time steps for the left hippocampus. The weighted degree-estimated *a* pair values are averaged across all subjects within the study dataset.

Therefore, when a high-weight interaction is detected at a time step *t* between two nodes with overall low temporal weight levels, that tie is considered as significant. Moreover, high correlation exists between the MLE based estimated values of distribution means (*a_i_.a_j_*) and the degree of the nodes, calculated as the sum of weights of the edges over *τ* time intervals. An example of such correlation is shown in figure 2 for left hippocampus (283 voxels) (further empirical results are provided in the supplementary information, figures 1–4) where the aggregated node degree-estimated latent variable *a* were averaged across all subjects of the study data. Also, as figure3 shows, there exists a weak negative correlation between the share of significant ties that is connected to each node *i*, and the MLE estimated variables *a_i_* corresponding to it. The share of significant ties are calculated as the number of ties connected to node *i* that are admitted to the final backbone network divided by the total edges connected to it (*N* – 1). The results of this analysis on other regions are provided in the supplementary information. These result establish the property that, thanks to the temporal null model, the admissibility of an edge *w_i,j_* into the irreducible network is not attributed merely to its degree, therefore controlling for the effect of local strengths of a node on the admissibility of its links to the backbone network.

**Figure 3:**
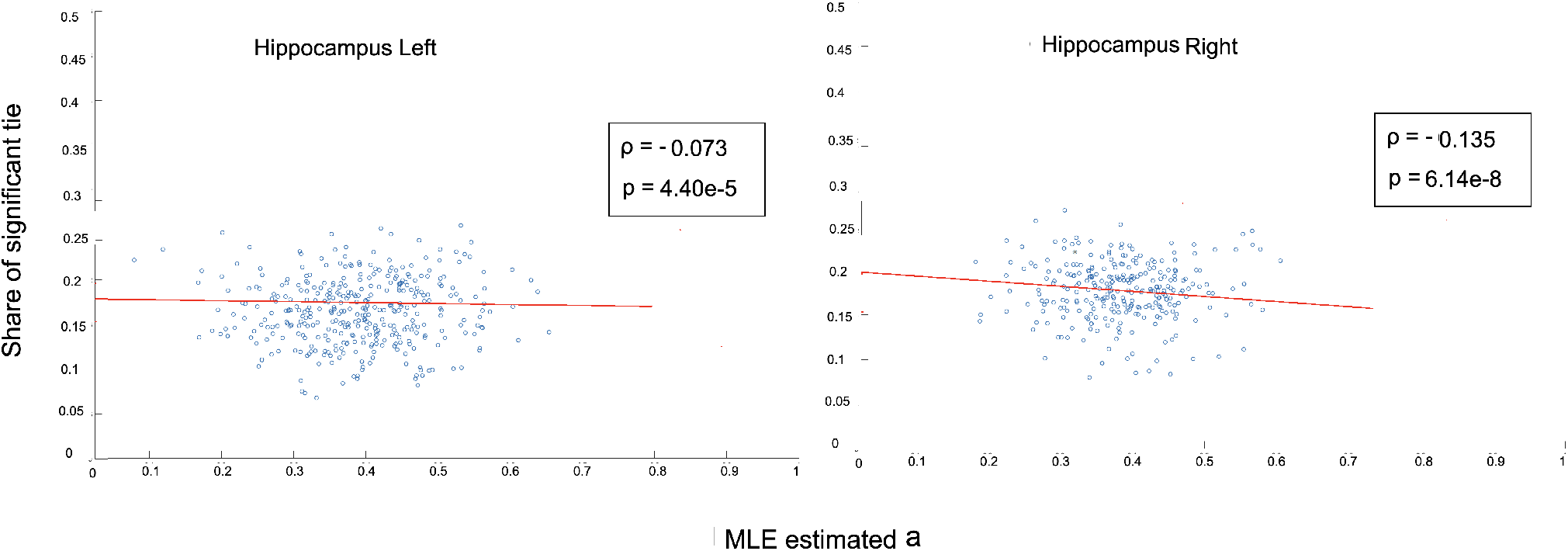
Correlation between share of significant ties connected to each node and the MLE estimated latent variable *a* for right and left hippocamous regions.

**Figure 4:**
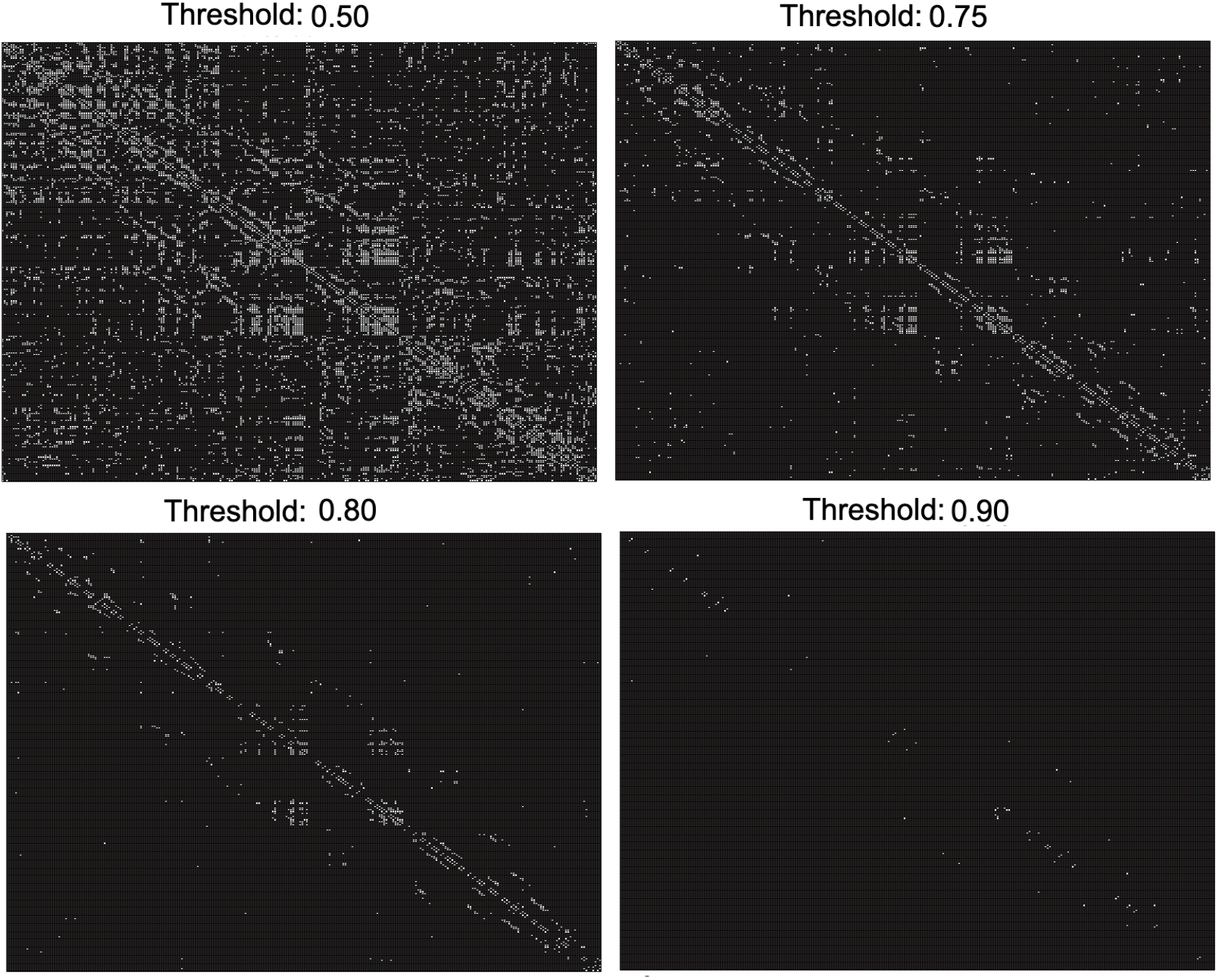
Derived backbone networks of the right hippocampus from one control subject given four threshold values.

## 4. Experimental results

In order to assess the proposed methodological framework, we apply it to a resting state fMRI data set of 300 subjects from the Autism Brain Imaging Data Exchange (ABIDE) database, including 150 subjects diagnosed with Autism Spectrum Disorder (ASD) (Di Martino et al. 2014). This dataset was selected from the C-PAC preprocessing pipeline and was slice time and motion corrected, and the voxel intensity was normalized using global signal regression. The automated anatomical labeling atlas (AAL) was then used for parcellation of regions of interest (Tzourio-Mazoyer et al. 2002). Then, the temporal links between each pair of nodes were extracted based on the Pearson correlation between their BOLD activation time series within each temporal window *t*, and were then rescaled based on min-max feature scaling to have continuous values within the range [0, 1]. The implementation code for the methodology in this work is available in https://github.com/ThisIsNima/Weighted-Backbone-Network.

After extracting the backbone networks on the referred dataset, we probed several aspects and measures of them, including the number of their significant ties and its relation with the *α* threshold, the structural comparison between the irreducible networks across various thresholds and neurological cohorts, the effect of temporal segmentation window size on the inferred networks, the relation between the weights of dFC ties and the inferred significant ties, and the performance of this approach on detecting injected random ties. For this analysis, we provide a closer assessment of backbone networks on four brain regions, namely the left and right hippocampus and the left and right amygdalas. We also provide part of the experimental results for the cerebellar regions in the main manuscript and the rest in the in the supplementary document. The reason for choosing these regions is the extensive focus of prior literature related to diagnosis and pattern discovery in functional connectivity among ASD patients on them and the fact that several types of abnormality have been discovered related to these regions among this group of patients (Shen et al. 2016; Cooper et al. 2017; Ramos et al. 2019; Guo et al. 2016; Rausch et al. 2016).

As the first step of our analysis, we examined the normality of the distribution of temporal links between each pair of nodes *i, j* across the experiment time *τ*. For this purpose, we used the Kolmogorov-Smirnov test on temporal ties between each pair of nodes for four different window sizes Δ ∈ {5, 10, 15, 20}. Table 1 demonstrates the average *p* values of the normality tests for the distribution of temporal ties between every pair of voxels across 300 subjects for four separate regions. These results demonstrate that the *p* values are below the 0.005 common threshold for rejecting the null hypothesis. Furthermore, the *p* value tends to increase as the size of temporal windows decreases, which can be attributed to the increase in total number of temporal windows *τ*. Beyond the theoretical basis of Central Limit Theorem, these results further highlight that the assumption of normality for the distribution of temporal edges in our resting state fRMI data is reasonable.

**Table 1:** p values of Kolmogorov-Smirnov test for normality of the distribution of temporal links. The p values presented in the table are averaged across all links ((*N*(*N* – 1))/2 edges for *N* nodes) of the network.

| Brain region | $\Delta = 5$ | $\Delta = 10$ | $\Delta = 15$ | $\Delta = 20$ |
| --- | --- | --- | --- | --- |
| L hippocampus | 2.1004e-11 | 2.4801e-9 | 1.1422e-22 | 2.5488e-32 |
| R hippocampus | 8.1161e-13 | 2.5102e-10 | 1.1084e-9 | 1.1100e-9 |
| L Amygdala | 6.3835e-14 | 1.3560e-9 | 1.1609e-9 | 1.1102e-9 |
| R Amygdala | 3.3875e-10 | 5.0045e-7 | 1.4108e-7 | 3.0545e-7 |

The backbone networks of the right hippocampus (region 38 per AAL atlas) for one control subject based on four different threhsolds (*c* = 1 – *α*) are provided as heatmaps in figure 4, where each cell represents a voxel, and white cells represents the significant ties. Note that self links are removed from these networks, thus the value of the diagonals of the heatmaps are set to zero. For this analysis, time courses were segmented into 20 temporal windows through the sliding window approach, with an overlap of 5 time points between consecutive windows (this is the default setup for the other parts of the experiments. Otherwise, we mention the temporal window size set up). The visualizations in figure 4 indicate that the number of admitted links decreases by increasing the threshold *c*. Moreover, the links between voxels in the vicinity of the diagonal line tend to endure the increase in threshold *c*, which highlights the strength of links between spatially close voxels.

The relation between the threshold *c* and number of significant ties is further inspected in figure 5-c, where the threshold increases from 0.5 to 1 with a fixed step resolution of 10^-2^. The number of significant ties for the network within each time window *t* = 1,…, *τ* is also provided in figure 5 a and b, where the red bars show the number of edges admitted to the final backbone network. As noted earlier, only the ties that meet the significance threshold in over 50% of the time steps *τ* qualify to be included in the final backbone network (red bar), thus the number of final admitted links is usually smaller than the significant ties within various temporal windows. However, the number of significant ties does not demonstrate a large variation across different temporal windows.

**Figure 5:**
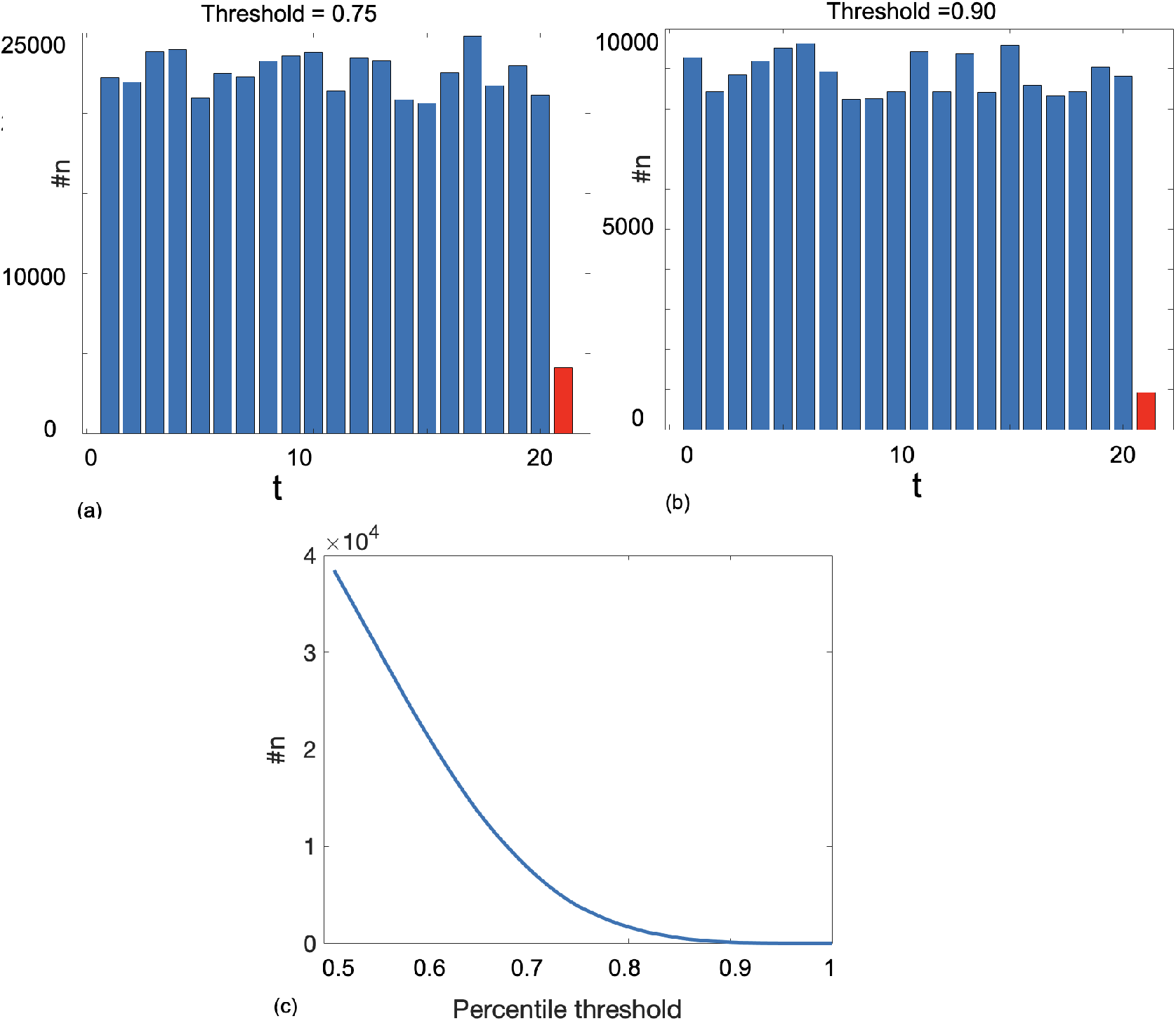
figures a and b: number of significant ties *n* for the right hippocampus network of one control subject across *t* = 1,…, 20 time steps based on two different threshold values. The red bar corresponds to the number of edges of the temporal network admitted to the backbone network. figure c: the number of admitted edges to the final backbone network based on 50 different threshold values between 0.5 and 1 with fixed resolution.

For the next step of the analysis, we examined and compared the backbone networks of the two cohorts (control and ASD) within our experimental dataset with similar temporal segmentation as the previous step. For this analysis, the value of *α* was set to 0.2, i.e. c-th percentile = 0.80. Figures 6 and 7 present the networks of significant ties extracted from the dynamic connectivity of the left and right hippocampus from four subjects, including two subjects diagnosed with ASD, and figures 8 and 9 show the extracted significant ties from the left and right Amygdalas for eight subjects, four of whom were diagnosed with ASD. Moreover, in order to provide a more comprehensive perspective of the irreducible networks of the mentioned regions, the averaged backbone networks of the two cohorts (Control and ASD) across the entire dataset are presented in figures 10 and 11.

**Figure 6:**
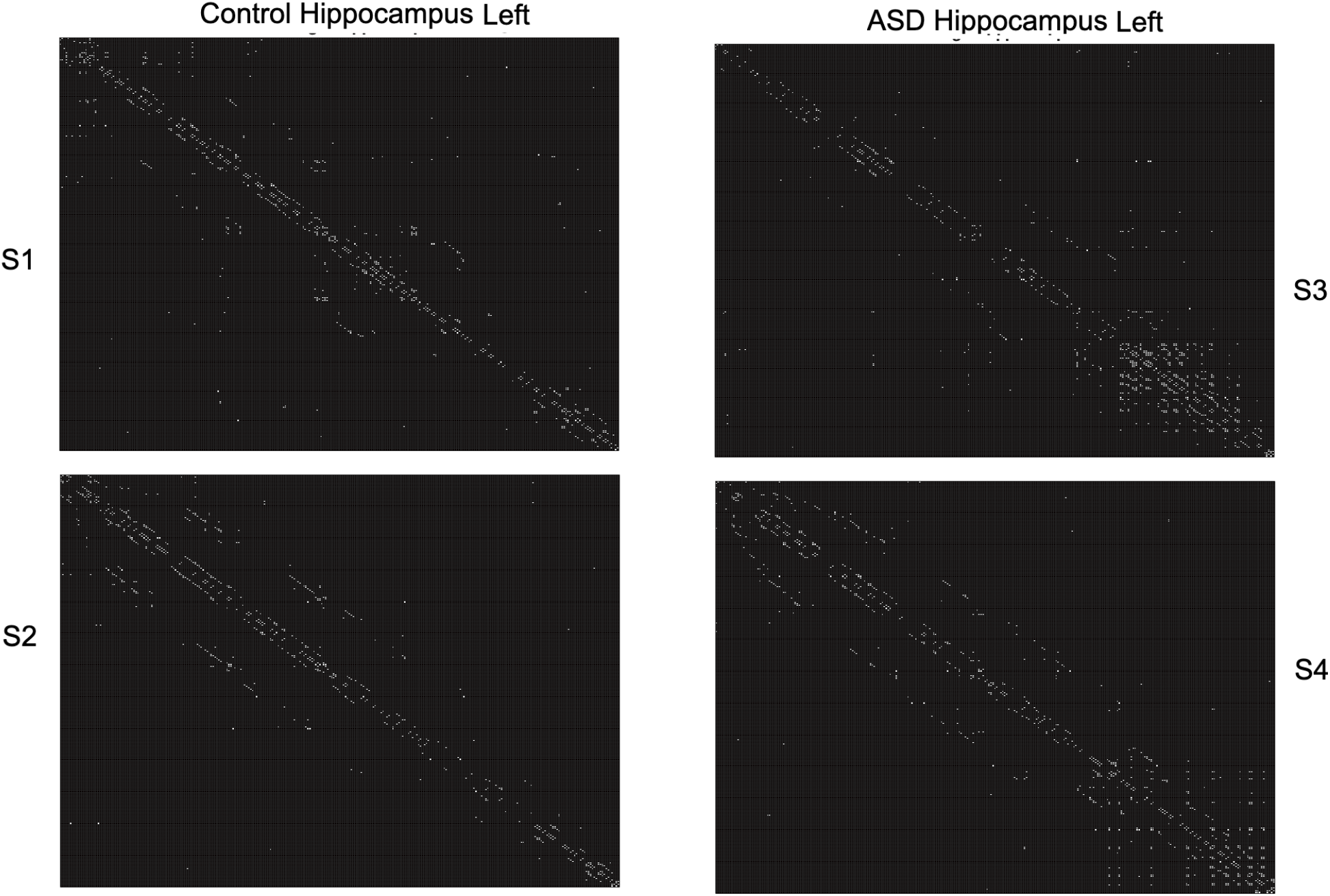
Extracted backbone network of the left hippocampus from four subjects; two control subjects and two diagnosed with ASD, based on a 0.80 threshold.

**Figure 7:**
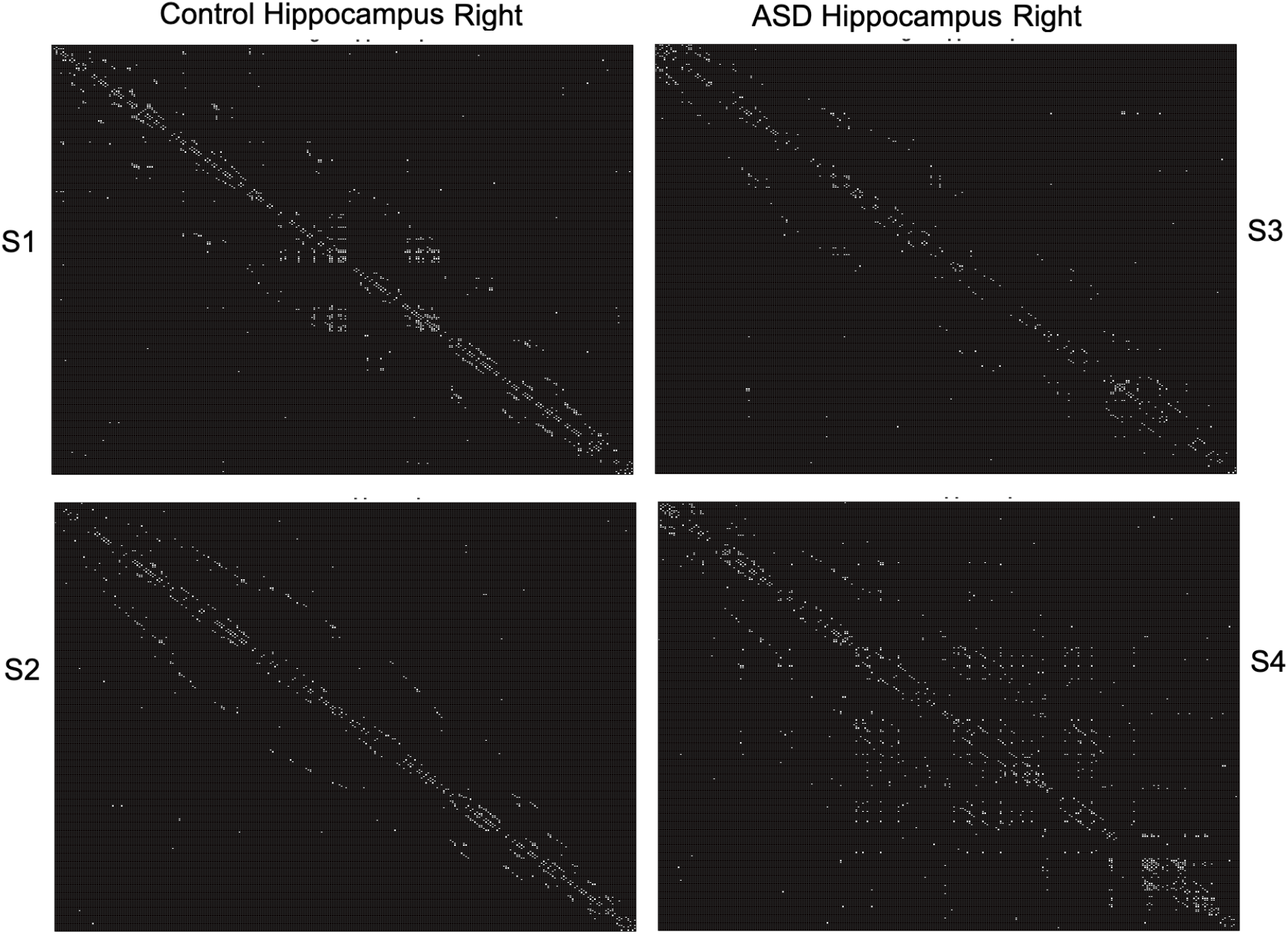
Extracted backbone network of the right hippocampus from four subjects; two control subjects and two diagnosed with ASD, based on a 0.80 threshold.

**Figure 8:**
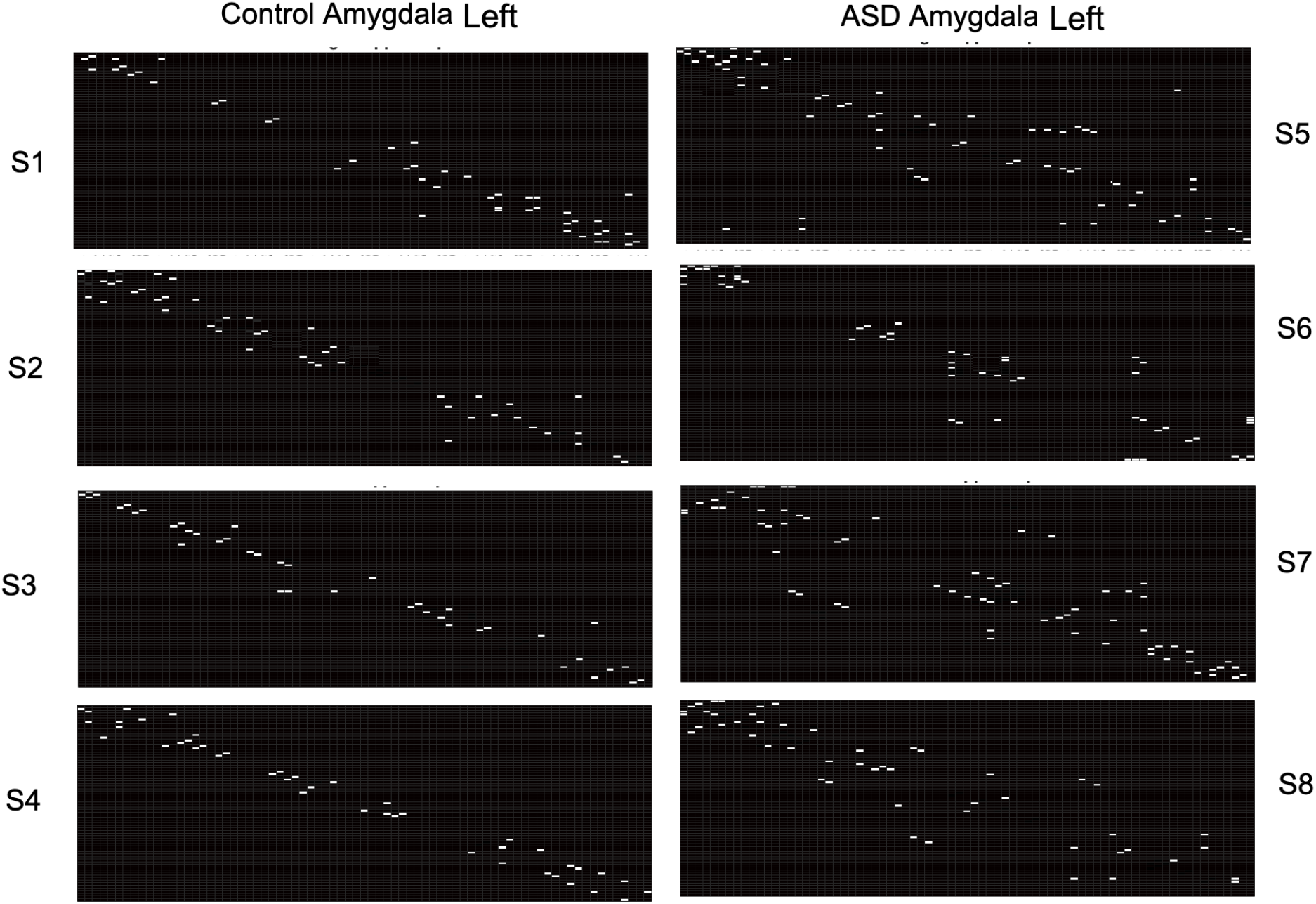
Extracted backbone network of the left Amygdala from eight subjects; four control subjects and four diagnosed with ASD, based on a 0.80 threshold.

**Figure 9:**
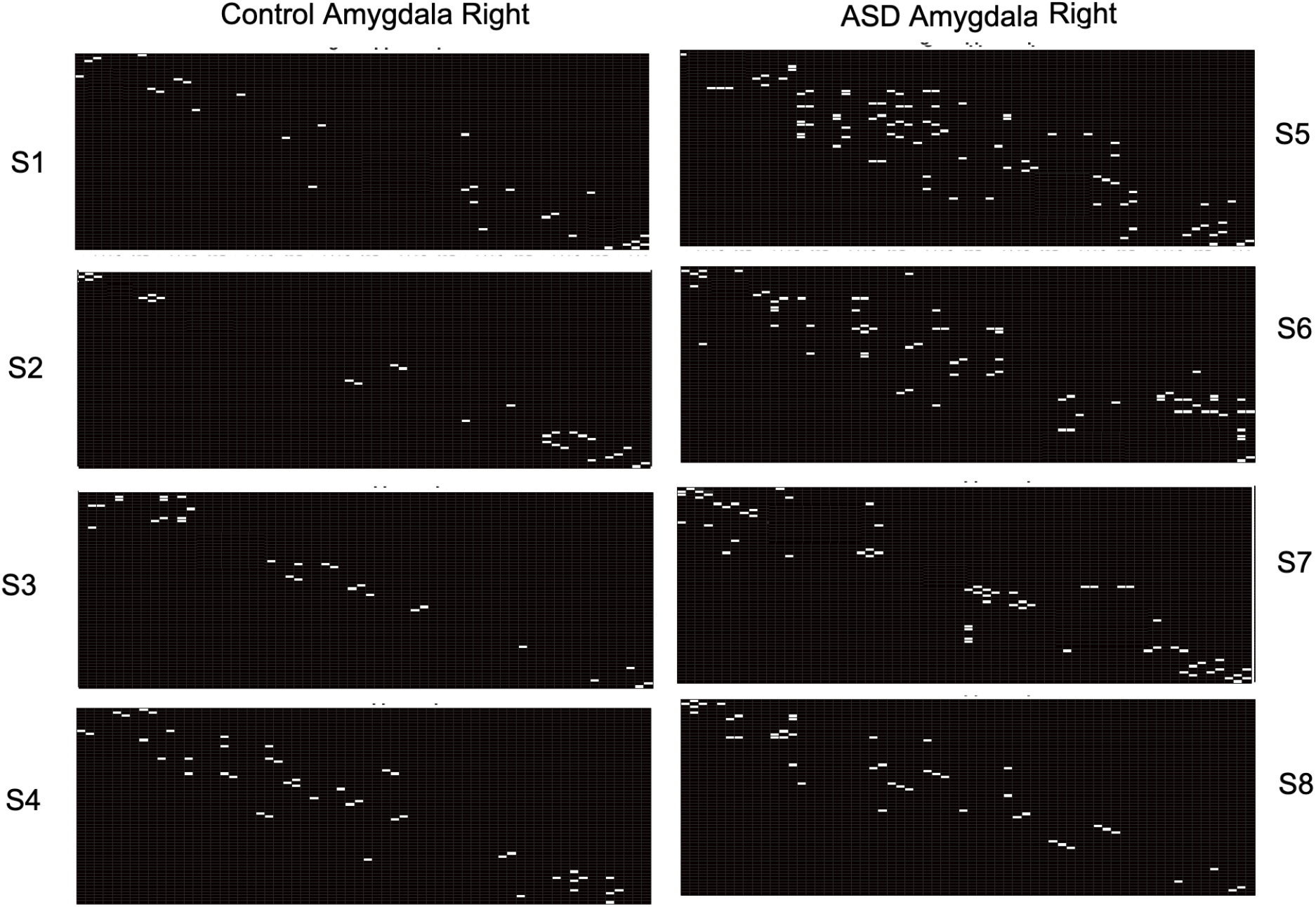
Extracted backbone network of the right Amygdala from eight subjects; four control subjects and four diagnosed with ASD, based on 0.80 threshold.

**Figure 10:**
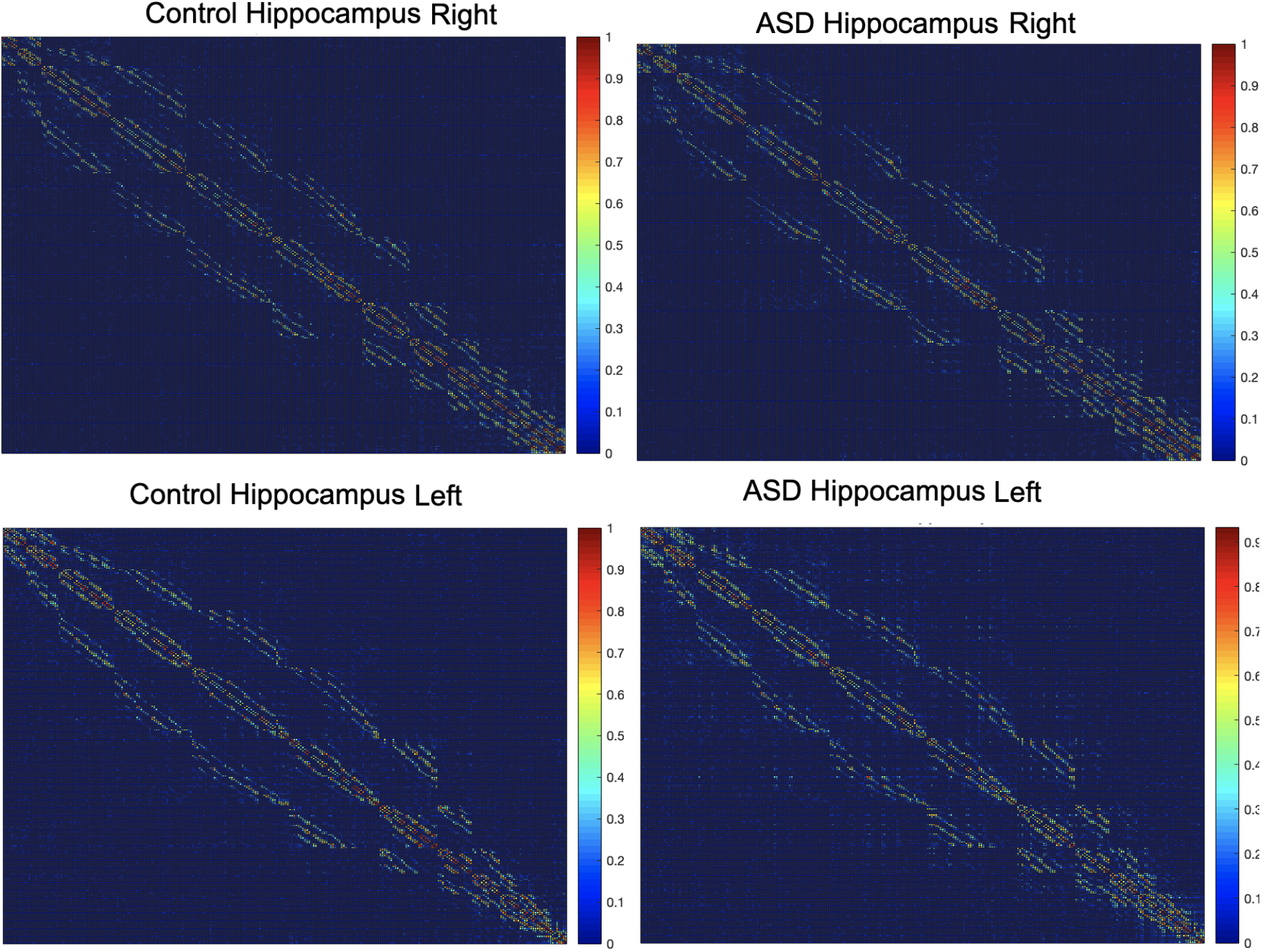
Average backbone network of the right and left hippocampus from 300 subjects; 150 control subjects and 150 diagnosed with ASD, based on a 0.80 threshold.

**Figure 11:**
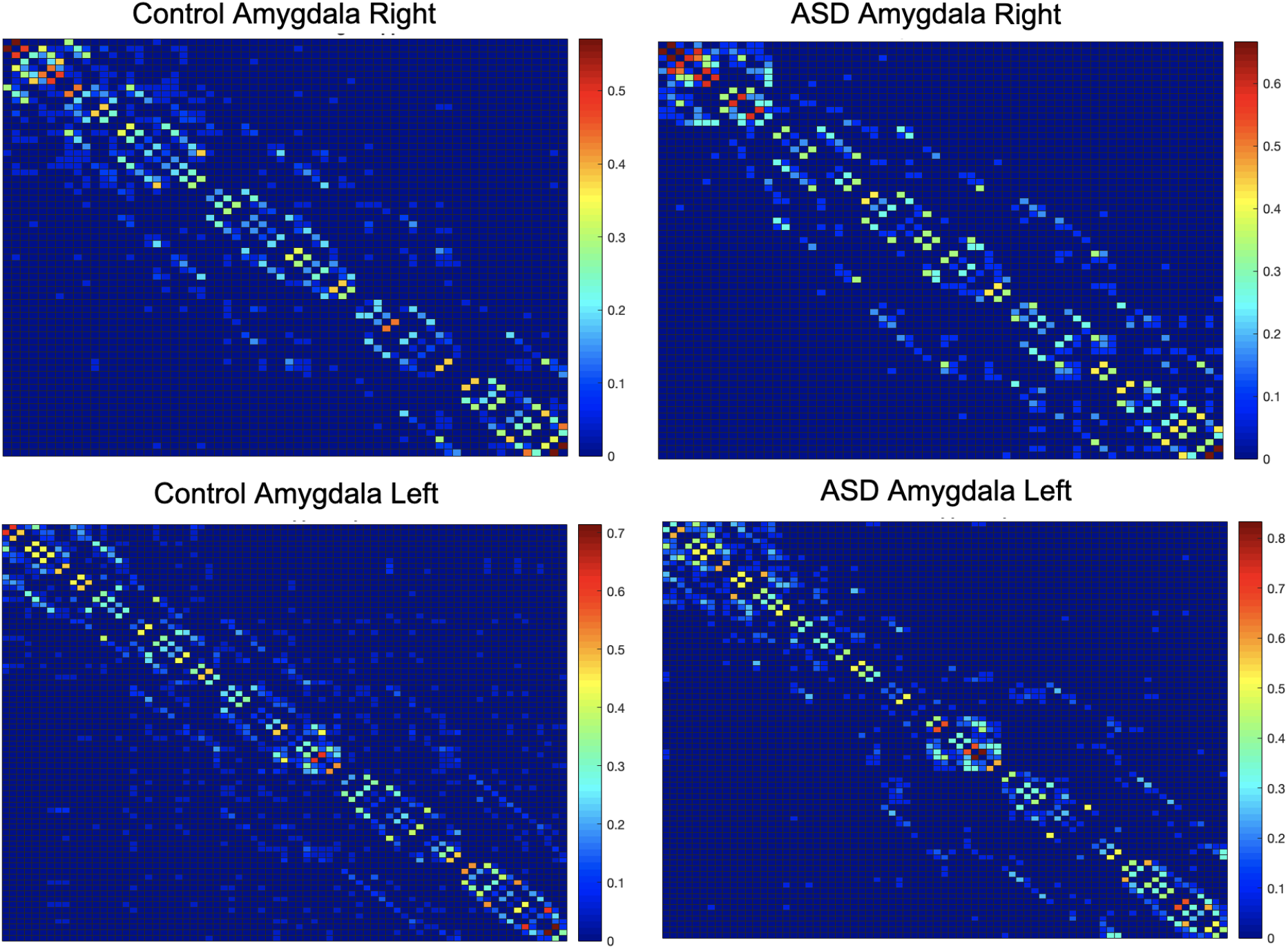
Average backbone network of the right and left Amygdalas from 300 subjects; 150 control subjects and 150 diagnosed with ASD, based on a 0.80 threshold.

Furthermore, we assessed the effect of the size of temporal windows on the extracted significant ties. For this purpose, we measured the difference between backbone networks of dFC based on four different window sizes: Δ ∈ {5,10,15, 20}, where the overlap between consecutive windows was 2 time points for the smallest window (Δ = 5), and 5 time points for the other three window sizes. As the measurement of dissimilarity, we used the mean percentage error (MPE) of the voxel-wise difference (between the values of corresponding matrix cells) between the backbone networks averaged across 300 subjects. The results of this analysis are provided in Figure 12 for two threshold values of 0.5 and 0.9. As this analysis demonstrates, the dissimilarity between the extracted backbone networks calculated as MPE is negligibly small for both temporal resolutions, which indicates the consistency of the backbone network against variations of the temporal window size.

**Figure 12:**
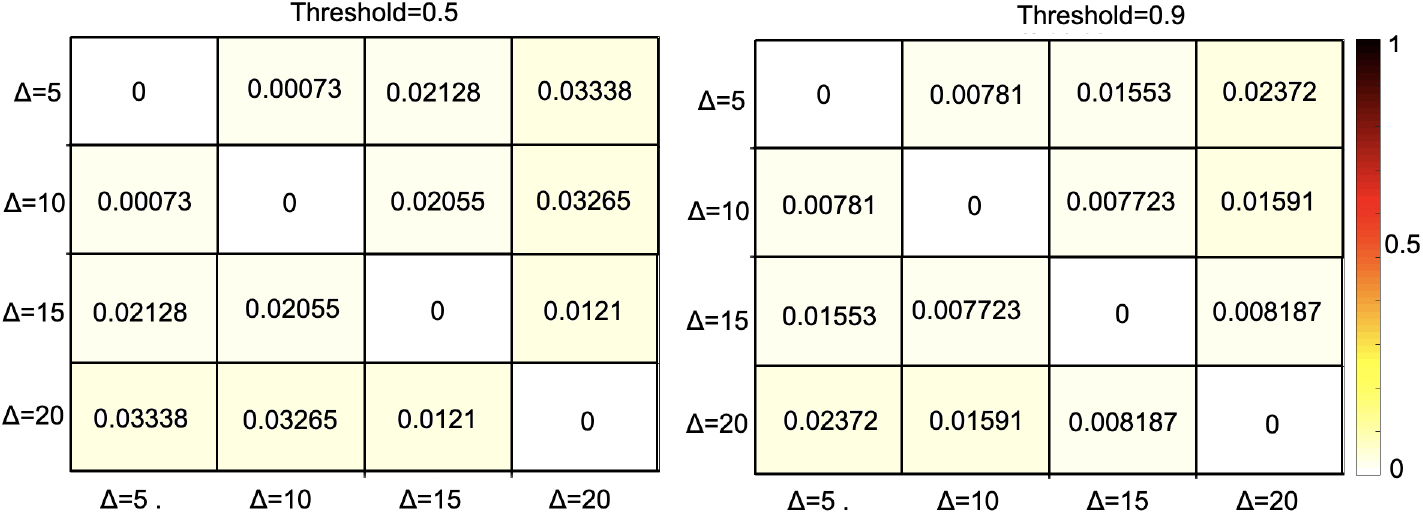
A comparison based on different window sizes using mean percentage error (MPE) of the voxel-wise difference between the backbone networks of dFC averaged across 300 subjects

As the last part of the voxel-level experiments, we examined the correlation between the empirical weight of the links and degree of the nodes in dynamic functional connectivity network with the backbone link wights and estimated latent variables *a, b*. In figures 13 a and b, the average backbone network of the right amygdala of 300 subjects as well as their average dFC over *τ* windows are presented. Additionally, the correlations between node degrees of the dFC network, calculated as the sum of the weights of temporal links for each node, and their estimated *a, b* based on MLE procedure, and the correlation between the average backbone link weights of 300 subjects and the average weight of their corresponding dFC links over *τ* windows is provided in Figure 13 c. Experimental results for additional regions of interest are provided in supplementary information. As these result demonstrate, there is a weak correlation between the weight of the dFC links and the average weight of backbone links (note that averaging binary backbone links results in continuous weights). Additionally, there is a relatively small negative correlation between node degree of dynamic functional connectivity and estimated distribution latent variables *a, b*. In line with the argument provided in the methodology, these empirical results further illustrate that WBN considers global and temporal information of the network beyond the local node degree and the weight of the links in the dFC network for extracting the significant temporal interactions.

**Figure 13:**
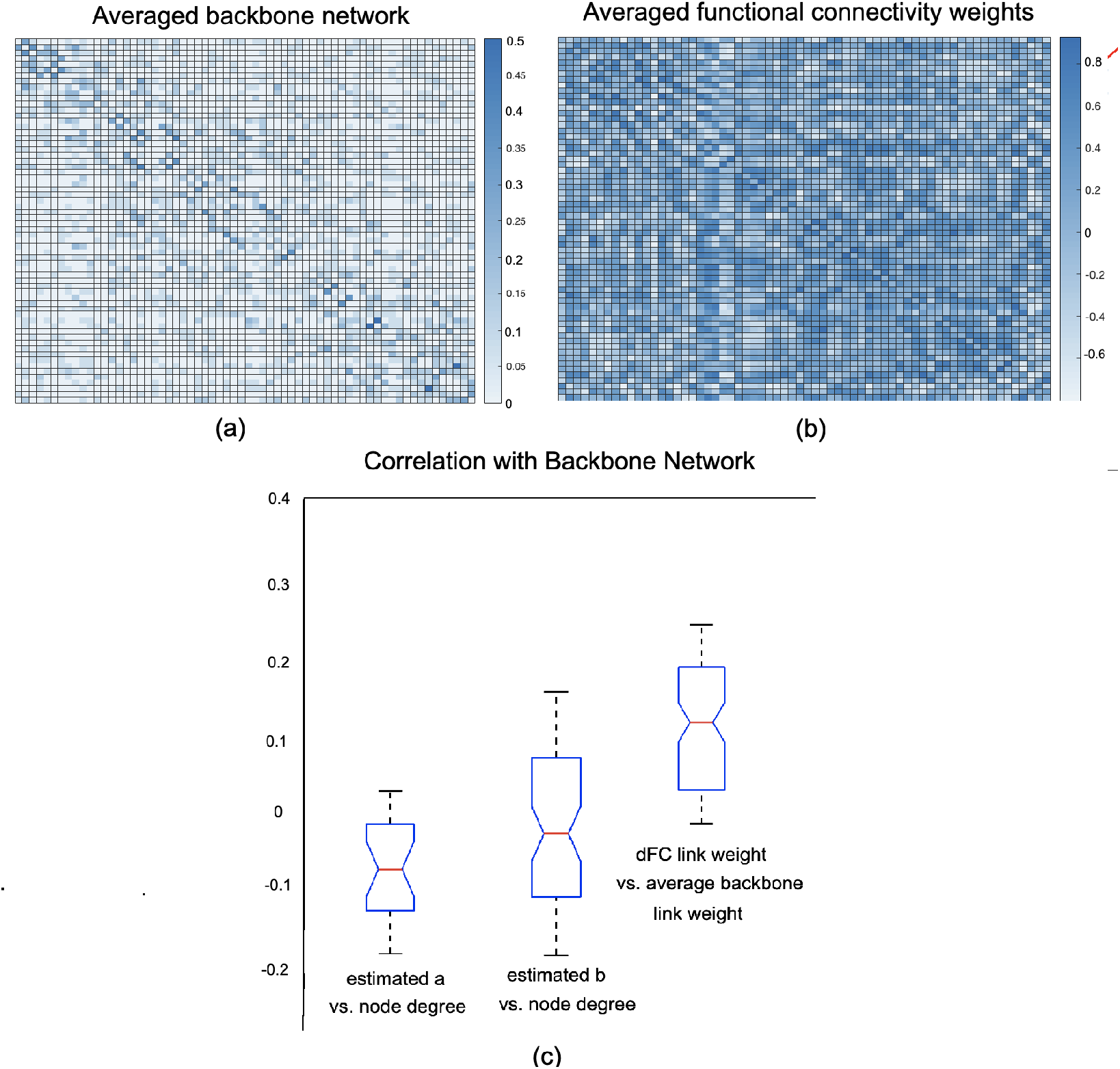
Figure a: average backbone network of the right Amygdala for 300 subjects. Figure b: average dFC network of the same region for the same subjects across *τ* = 20 time intervals. Figure c: correlations between average node degree and estimated *a, b* backbone link weights as well as average dFC link weight over *τ* = 20 intervals for 300 subjects.

### 4.1. Full brain Analysis

Just like the voxel-level analysis, a full brain backbone network of rsFC can be extracted where each node is a region of interest (ROI). For this purpose, the time courses within each region based on the AAL atlas was averaged. The averaged full-brain networks of significant ties for 150 control and 150 ASD subjects are demonstrated in figures 14, 15, 16, and 17, where the seed regions are the left and right amygdalas and hippocampus, and the average backbone links with weights below 0.05 where filtered out to facilitate easier presentation. In these figures, the link weights (link thickness) correspond to the count of their corresponding links appearing in the binary backbone networks across each cohort. We can observe from the figures 16, and 17 that the amygdalas and hippocmapus develop a larger number of significant ties with other regions among the control group compared to the ASD cohort as the network of the latter cohort is more sparse. The width of links in figures 14 and 15 also represent the strength of the average correlations. We can therefore also observe a relatively high averaged backbone link between the right and left hippocampus among both the control and ASD group. However, certain differences can be detected between the two cohorts, including a stronger average backbone tie between the hippocampus and the cerbellums as well as right and left olfactories among control subjects compared to the ASD cohort. Moreover, a higher average backbone tie can be noticed between the left and right amygdalas and the superior and the middle temporal gyrus among control subjects. Further related experimental results are provided in the supplementary information, which include the average backbone connectivity with several cerebellar regions being the seed area. These results demonstrate the benefit of the weighted temporal backbone network in revealing the differences in irreducible ties between different regions of interest.

**Figure 14:**
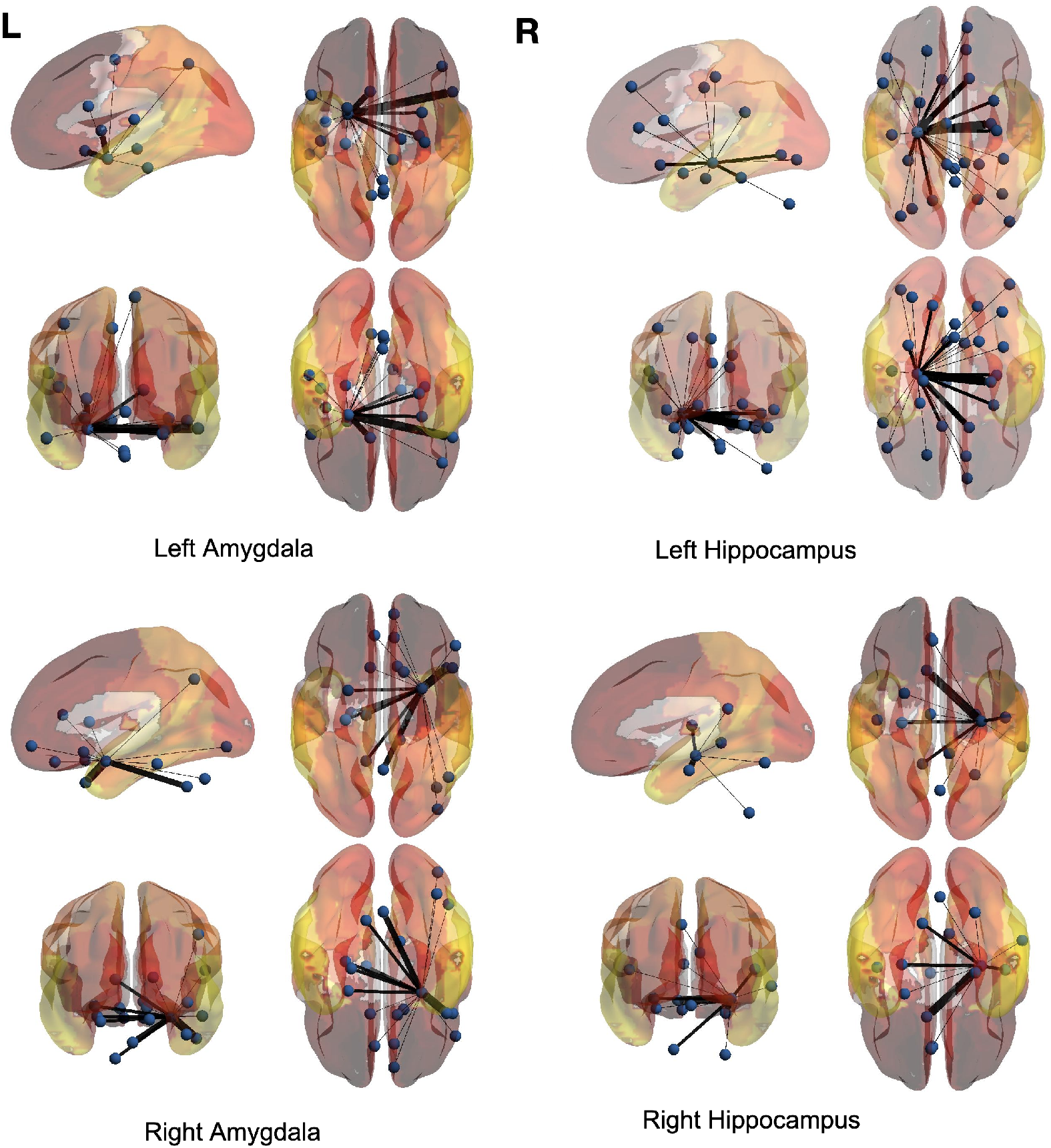
The averaged full brain backbone networks of 150 control subjects based on four seed regions

**Figure 15:**
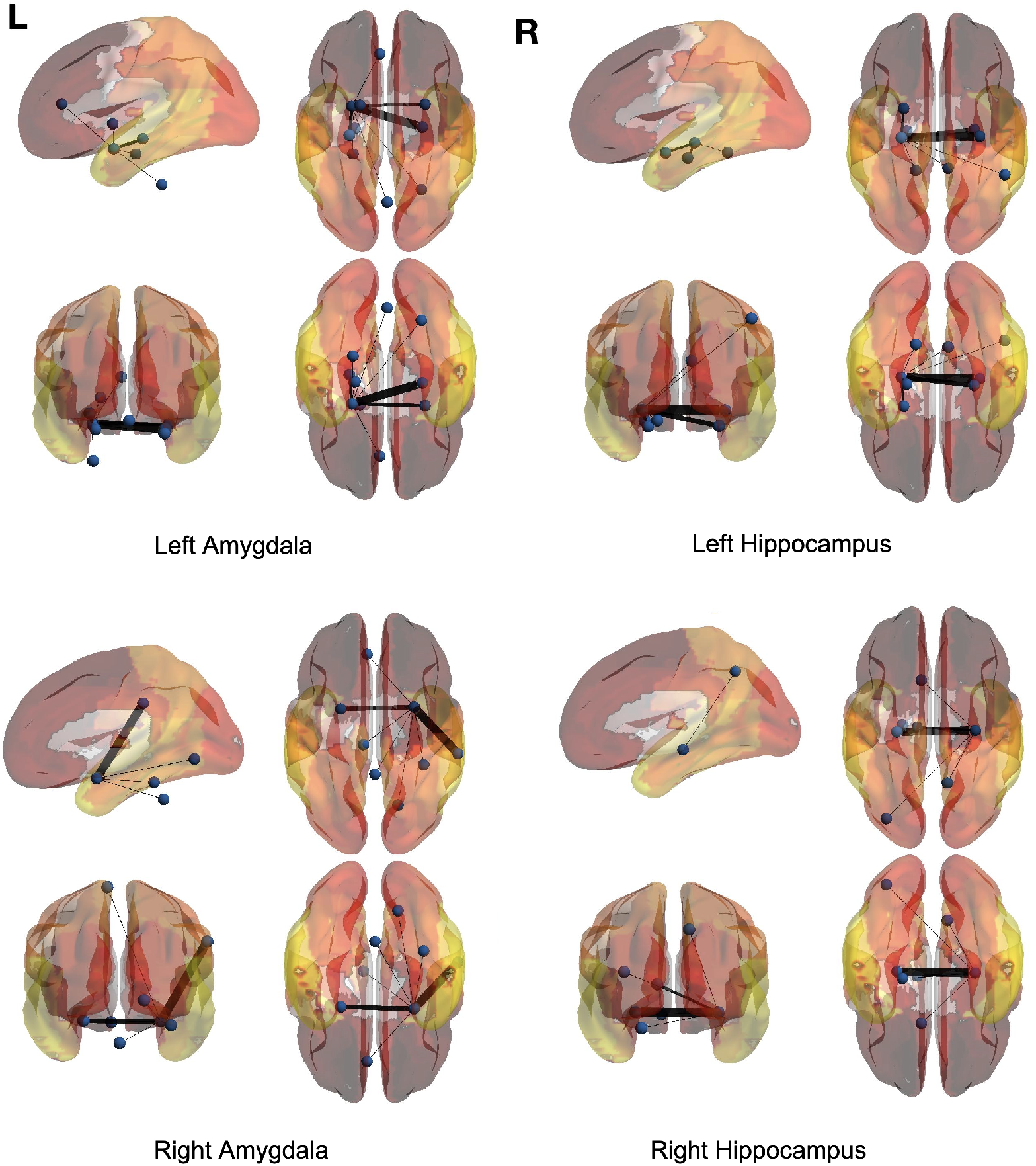
The averaged full brain backbone networks of 150 ASD subjects based on four seed regions

**Figure 16:**
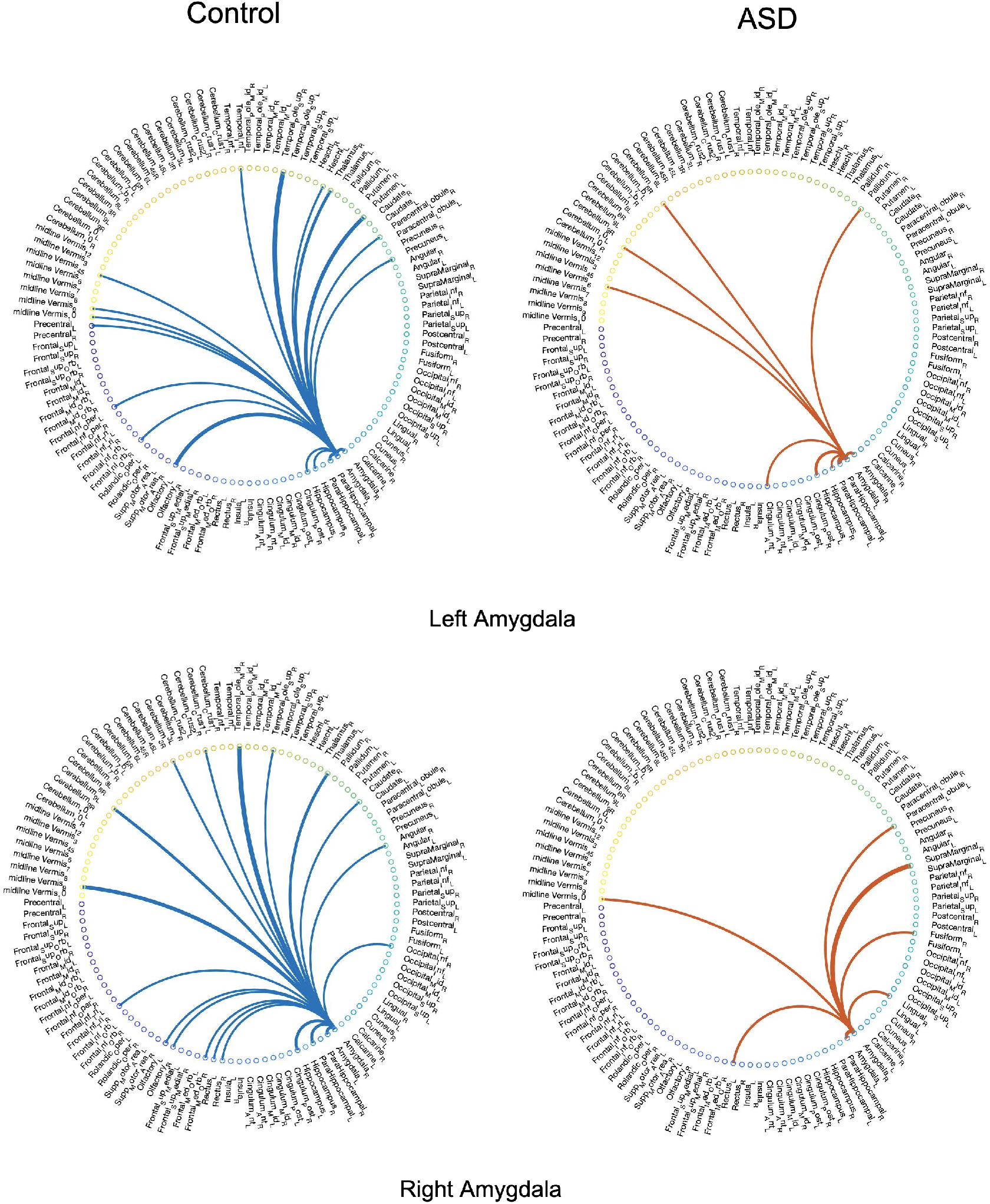
Averaged full brain backbone networks of 150 ASD and 150 control subjects with the left and right Amygdalas as the seed regions

**Figure 17:**
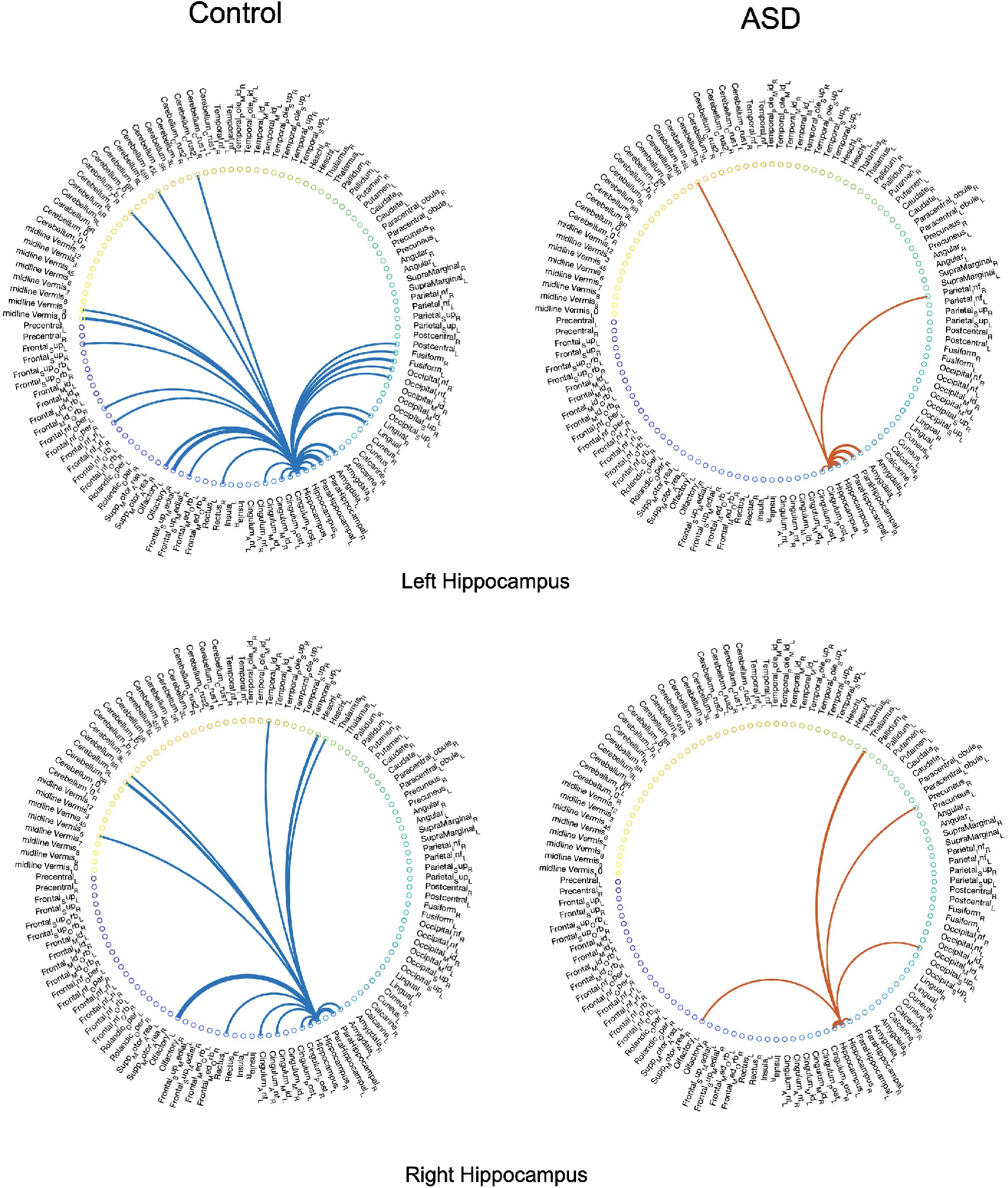
Averaged full brain backbone networks of 150 ASD and 150 control subjects with the left and right hippocampus as the seed regions

Also, figure 18 depicts the averaged backbone connectivity of the cerebellums (18 regions per AAL) and the vermis (8 regions per AAL) of the two cohorts in this study, which indicates higher connectivity level among the control group compared to the ASD group. These results, along with the experimental results provided in supplementary information (figures 5–10), can indicate that the increased cerebro-cerebellar functional connectivity detected in some studies can be driven by a large number of links that fail to be incompatible with the null hypothesis of links being produced at random. In other words, despite the lower connectivity detected in cerebro-cerebellar subnetwork among the control group in terms of number of links or their weights, the number of meaningful and irreducible links in that subnetwork among the healthy cohort tend to be larger compared to the ASD cohort (Ramos et al. 2019; Mostofsky et al. 2009; Khan et al. 2015).

**Figure 18:**
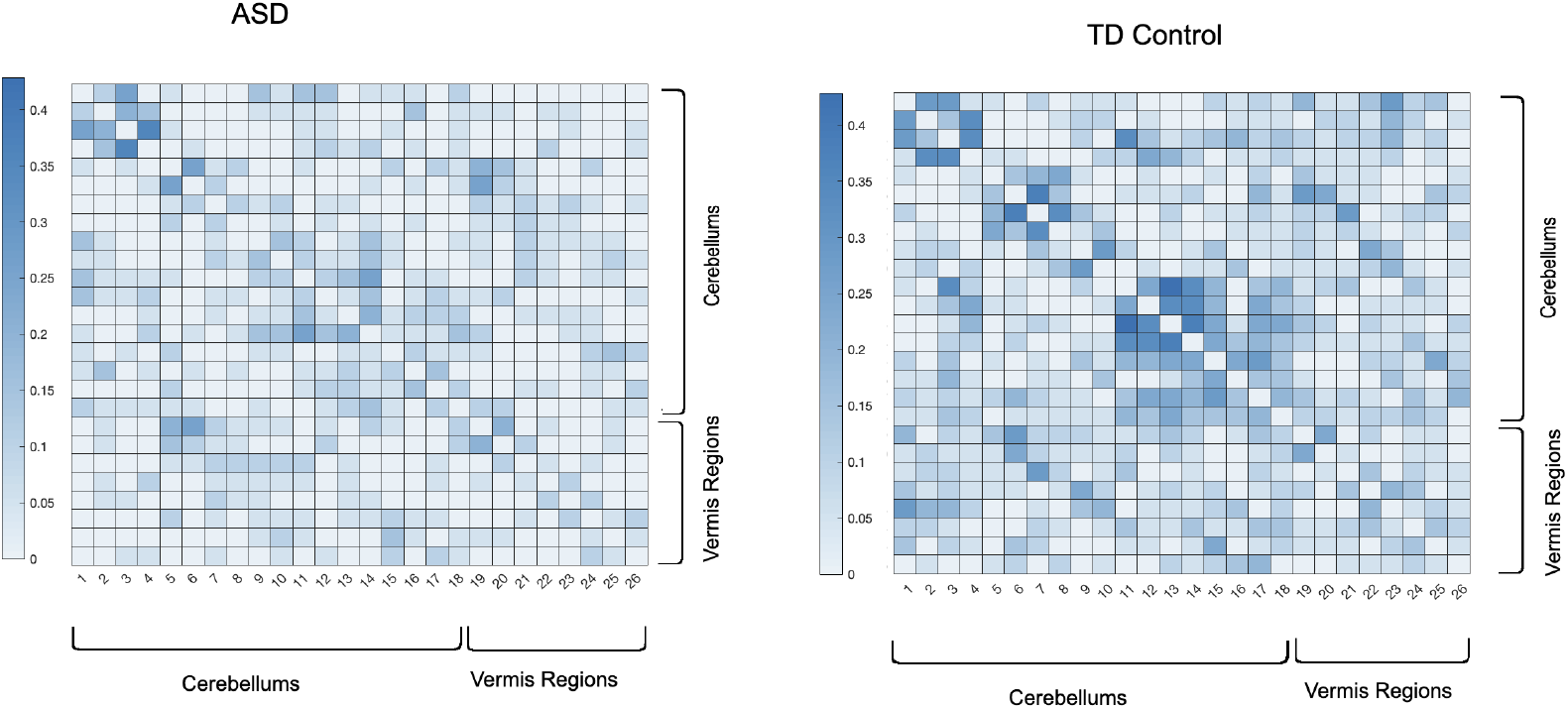
Averaged full brain backbone networks of cerebro-cerebllar and vermis regions across 150 ASD and 150 control subjects

### 4.2. Detection of random links

To further evaluate the performance of the proposed approach in filtering out non-significant temporal ties compared to the binary ST filter, we assigned random weights to a subset of edges of the real rsFC networks of our dataset. For this purpose, 100 random weights were injected into 100 links of rsFC of the left hippocampus, and the precision of WBN as well as the ST filter approach, proposed by Kobayashi et al., in excluding them from the final network were calculated (Kobayashi, Takaguchi, and Barrat 2019). Due to the fact that the ST filter operates on temporal binary weight networks, in order to evaluate it we converted the rsFC link weights as well as the randomly injected weights into binary links by drawing a temporal link between each pair of node whose weight in the original rsFC network was above the entire networkaverage. The result of this experiment is provided in figure 19, where WBN demonstrates an advantage over the ST filter in random link detection precision. Similar experiment with other regions of interest was conducted, which is provided in the supplementary information (figure 11). The evaluation measure for this analysis were calculated by comparing the detection of injected random link weights with the ground truth. Part of the superior performance of WBN can be attributed to the fact that the process of conversion to binary network for the ST filter setup results in loss of information and precision, which is an inherent disadvantage of backbone network detection approaches that are designed for binary networks.

**Figure 19:**
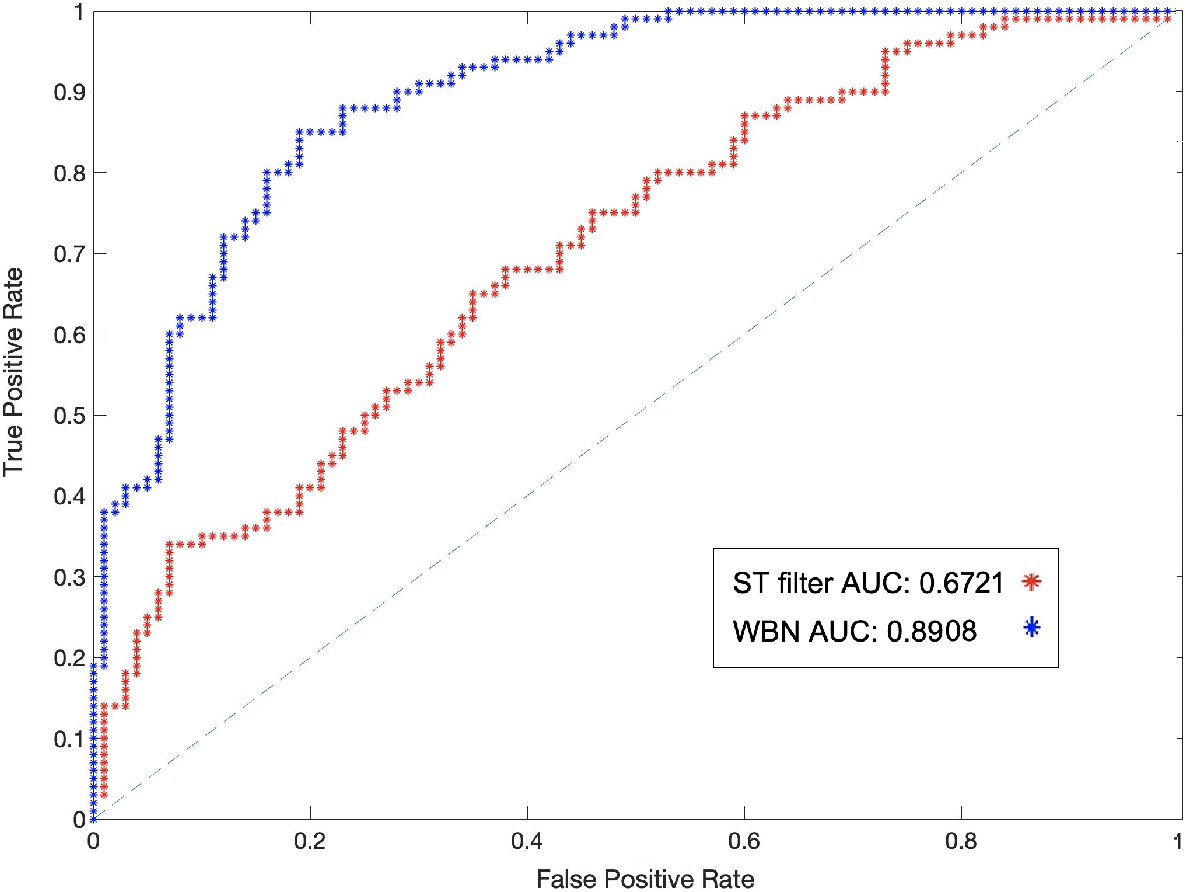
The AUC of detection of injected random weights based on the ST filtering as well as the proposed approach (WBN) in the left hippocampus.

## 5. Discussion

In this work, we proposed a new approach for detecting the significant ties between nodes on voxel and ROI level networks of resting state dFC. The proposed framework entails two computational steps; first, a maximum likelihood optimization calculates the pair of latent variables that drive the mean and standard deviation of the optimal Gaussian distribution of the temporal links between each pair of nodes across *τ* time steps. Then, the empirical link weights between each pair of node within each temporal window *t* ∈ {1, 2,…, *τ*}, denoted as 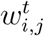, is compared to the *c–th* percentile of the Gaussian distribution characterized by the optimal mean and standard deviation calculated from the previous step, denoted as *w_c-th_*. If 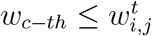, then we consider 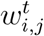 a significant tie, and if the temporal tie between *i,j* meets this significance threshold for more than %50 of the time over the *τ* windows, then the link between nodes i, j is admitted to the final backbone network of the resting state dFC. This process is performed for every pair of nodes in the temporal network of dFC in order to generate the backbone network. Aside from providing a systematic filtering framework for weighted temporal networks such as resting state dFC, this approach has several analytical advantages over other prior filtering approaches that we discuss in this section. We also discuss the limitations of the proposed methodology along with possible suggestions for improvement and future plans.

As mentioned previously, inclusion of a temporal link in the backbone network is determined by testing the hypothesis that the link can be explained by the null model that links are created uniformly at random. This comparison is applied to every link in the dFC individually, i.e. between every pair of nodes and within every temporal window. Therefore, temporal properties and variations of the network structure over time are taken into account in backbone network inference. This property is an advantage of the proposed methodology over some of the prior approaches that consider a constant intrinsic activity value for the nodes over time. It also offers the power of determining a cut-off percentage of ties having a larger weight over the *c – th* percentile, which was decided to be %50 in this study.

Another advantage of the suggested approach is the fact that it considers the interplay of global and local information of the network in estimating the latent variables *a* and *b*. In other words, the significance of temporal ties cannot be attributed merely to node properties such as degree or centrality measures, because each equation in the system of *N* equations of equations 4 and 5 takes into account the combination of weights over time for each link for node *i* as well as their combination with other links between *i* and every other node in the network. This property has been discussed in more detail in methodology section and evaluated in the results section.

The refinement of parameters of the distribution through maximum likelihood optimization requires solving the system of *N* equations for *N* nodes (one set of *N* equations for each of the two parameters), which can be solved through several optimization approaches such as gradient-based optimization, search methods, or the Newton method. Solving these equations does not require any hyper parameter tuning as the only parameters that need to be selected as input is the threshold value *α* and the percentage of times that the weight of the link meets the *c* = 1 – *α* percentile of the distribution, which offer the flexibility for a comprehensive assessment of the temporal ties in the dynamic connectivity network.

Unlike some of the null models suggested in the past for binary networks based on binomial or Poisson distributions, the methodology put forward in this work does not assume a strictly positive weight between interacting nodes. This property provides the flexibility for ties that are generated through various approaches such as correlation measures to be considered in the null model, as negative correlation is a possibility between interacting nodes.

Another advantage of the proposed approach is the fact that the backbone networks are learned for each subject individually. As explained in the methodology section, the input for WBN is the weighted dynamic connectivity network of a subject, and its output is the network of irreducible ties corresponding to the subject. This property has the benefit of taking into account the individual differences when inferring the backbone network in an isolated fashion.

The suggested methodological framework can be used in studies with various scales and resolution of dFC networks, meaning that instead of voxel-level analysis, dFC networks consisting of regions of various scales as nodes can benefit from this approach as well. Moreover, this approach is independent of temporal segmentation step, as long as the statistical properties of independence and normality are met.

### 5.1. Limitations

Despite the mentioned advantages, the proposed approach bears certain limitations which we highlight in this section.

As discussed in the methodology, the first step of the suggested framework entails estimation of latent variables *a* and *b*, which rule the propensities to generate a distribution of links with a certain average and standard deviation. However, these variables are estimated and compared across the experiment time *τ*, i.e. the length of the fMRI signals. In other words, the mean and standard deviation of the distributions, and in turn the backbone network calculations, can vary depending on the length of the experiment.

Another limitation of the suggested approach is the assumption of normality for larger temporal window sizes. As the empirical tests demonstrated, an increase in size of the temporal windows could in principle weaken the normality assumption of the distribution of the temporal links. Despite the evidence of normality for reasonable and common window sizes in the literature, this assumption needs to be further explored for various different datasets.

The MLE optimization for estimating the intrinsic variables *a, b* plays the largest role in the computational complexity of the methodology presented in this work. The computation time depends on the number of nodes, i.e. spatial resolution, and the number of time intervals that the signal is segmented to. By definition of the approach, the spatial resolution plays a more significant role in the computational complexity (refer to equations 4 and 5). In this study, the system of *N* equations were solved through the trust-region-dogleg method, whose computation time for regions below 1000 voxels was 10 minutes for 8 GB of RAM memory. However, more efficient approaches can be employed for this purpose.

Alleviating the mentioned limitations requires further methodological explorations and analytical studies on various datasets. As future work, our objective will include assessment of the backbone network of resting state dFC of other cohorts and data from various neurological conditions and to study different group differences. Furthermore, assessment of significant temporal structures and graph communities and motifs as well as exploring the effect of different preprocessing pipelines on the outcome of the proposed approach can be fruitful paths for further experiments in the area of dynamic functional connectivity.

## 6. Acknowledgement

This work was supported by National Institute of Health grants to I. R. Olson [R01HD099165; RO1 MH091113; R21 HD098509; and 2R56MH091113-11].

