## Supplementary Information for "The Backbone Network of Dynamic Functional Connectivity"

### Supplementary Material

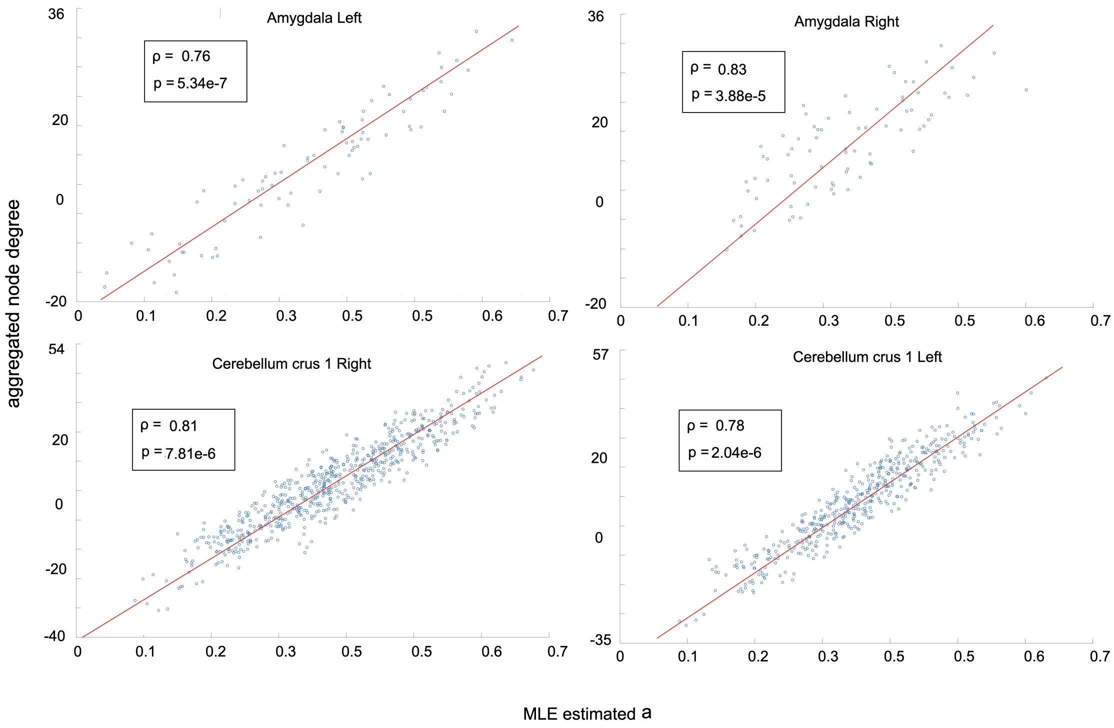


Figure 1: Correlation between node degree calculated as aggregated weights of all edges connected to each node over time $\tau$ and the MLE estimated latent distribution mean variable $a$

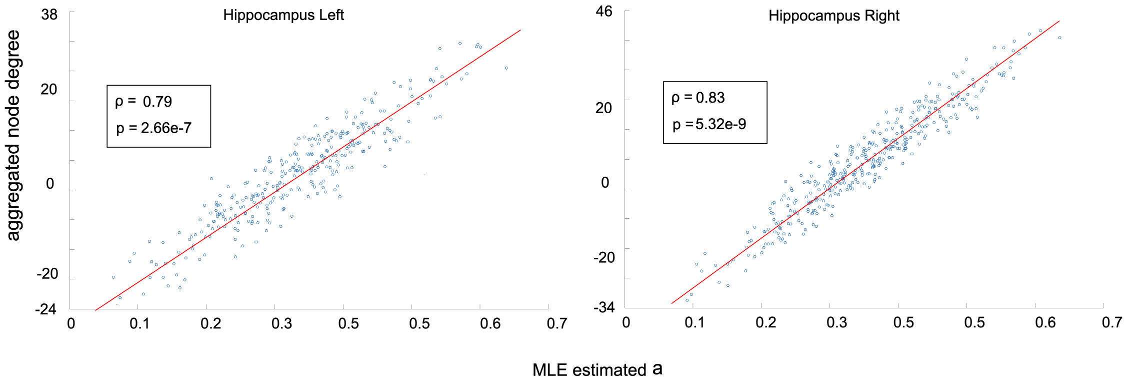


Figure 2: Correlation between node degree calculated as aggregated weights of all edges connected to each node over time $\tau$ and the MLE estimated latent distribution mean variable $a$

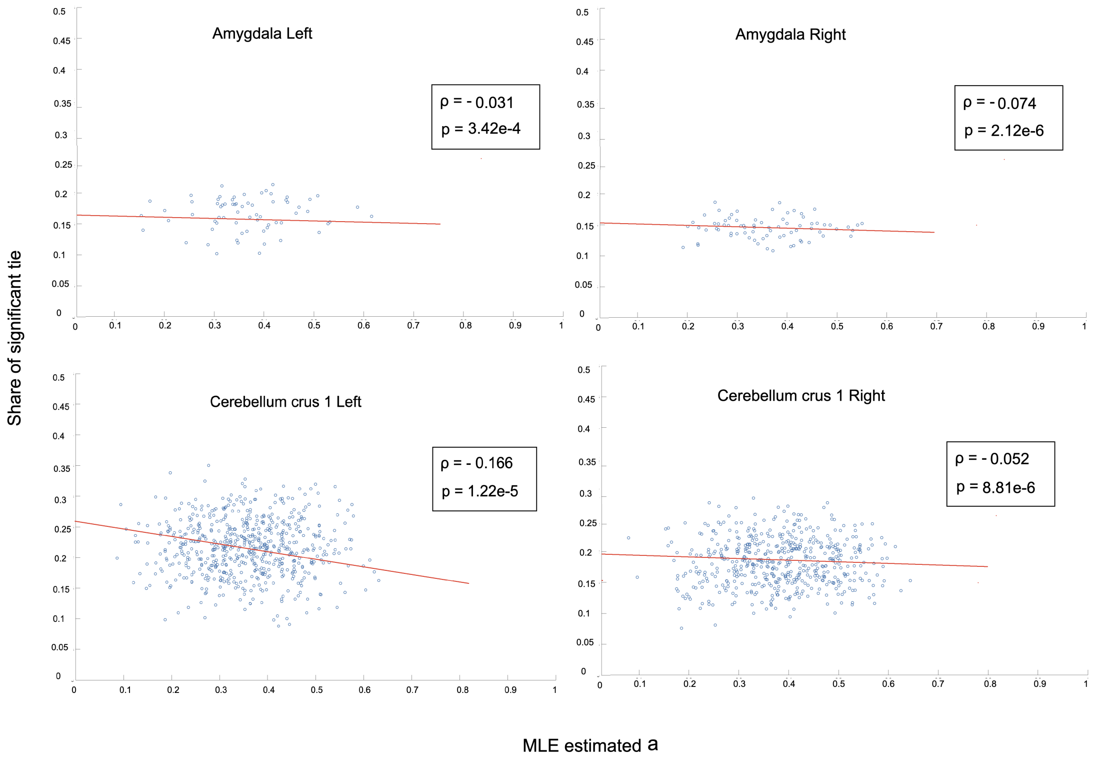


Figure 3: Correlation between the share of significant ties of each node over time $\tau$ and the MLE estimated latent distribution mean variable $a$

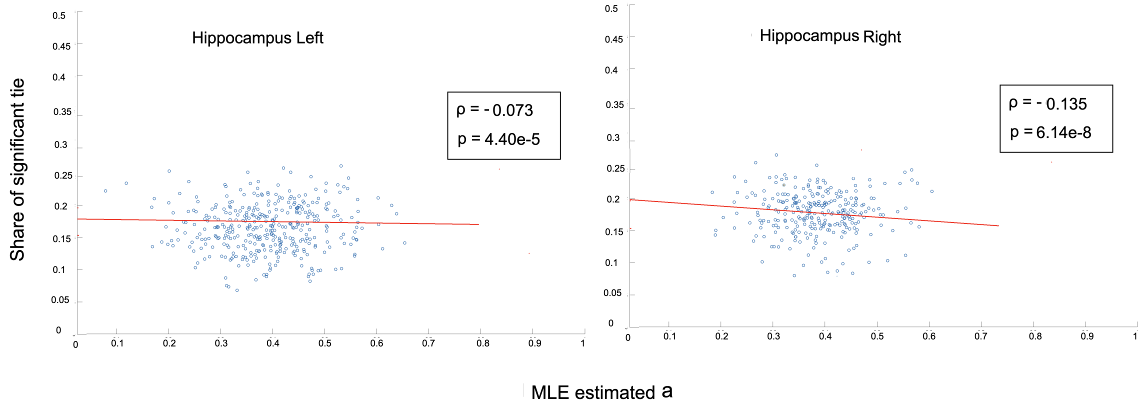


Figure 4: Correlation between the share of significant ties of each node over time $\tau$ and the MLE estimated latent distribution mean variable $a$

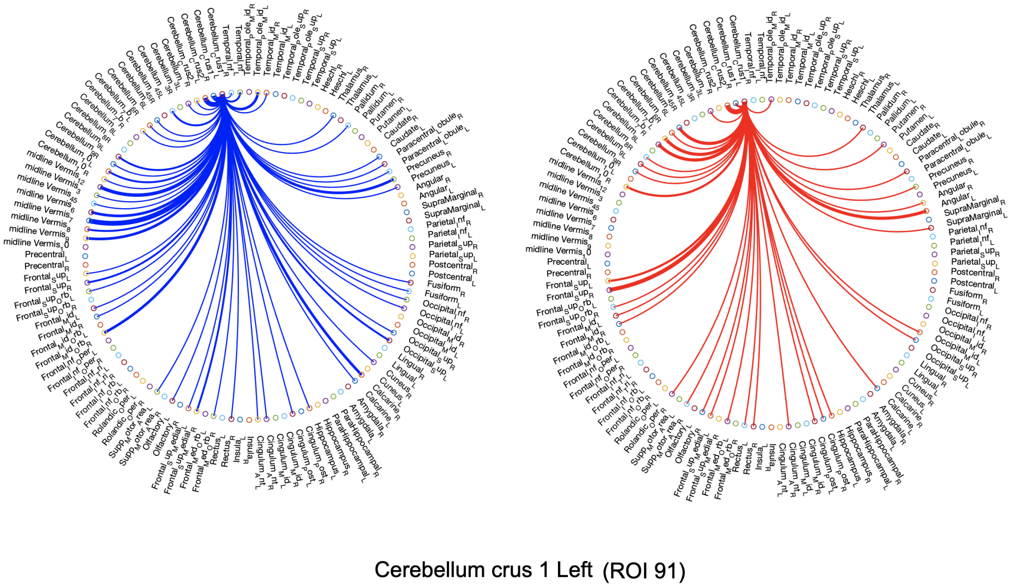


Figure 5: Comparison of average backbone connectivity between left cerebellum crus 1 and other regions between the control and ASD cohort

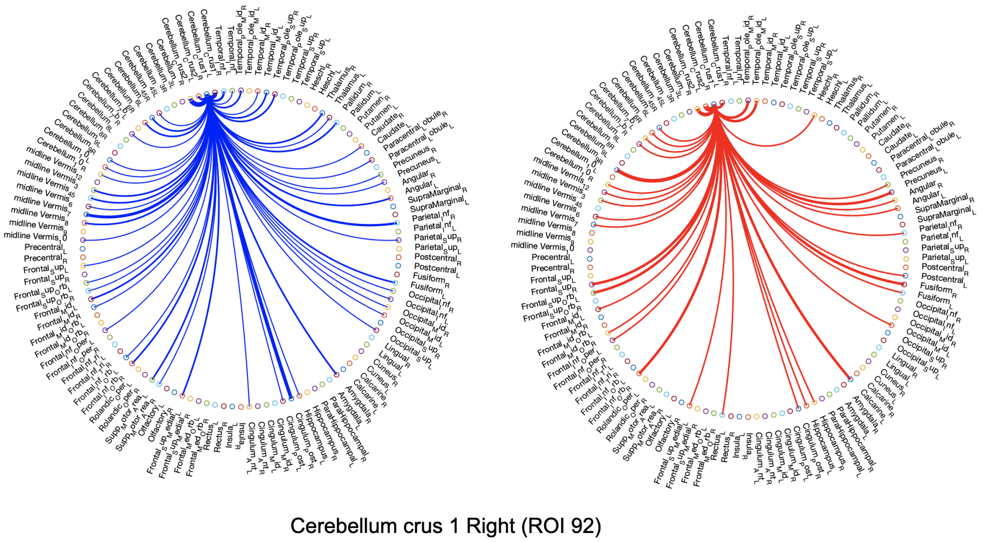


Figure 6: Comparison of average backbone connectivity between right cerebellum crus 1 and other regions between the control and ASD cohort

.png

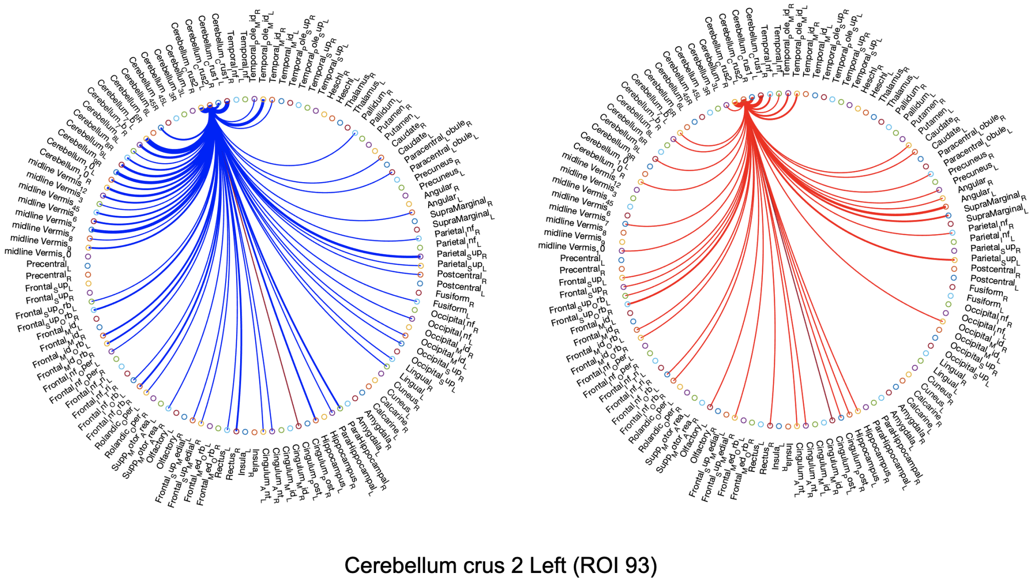


Figure 7: Comparison of average backbone connectivity between left cerebellum crus 2 and other regions between the control and ASD cohort

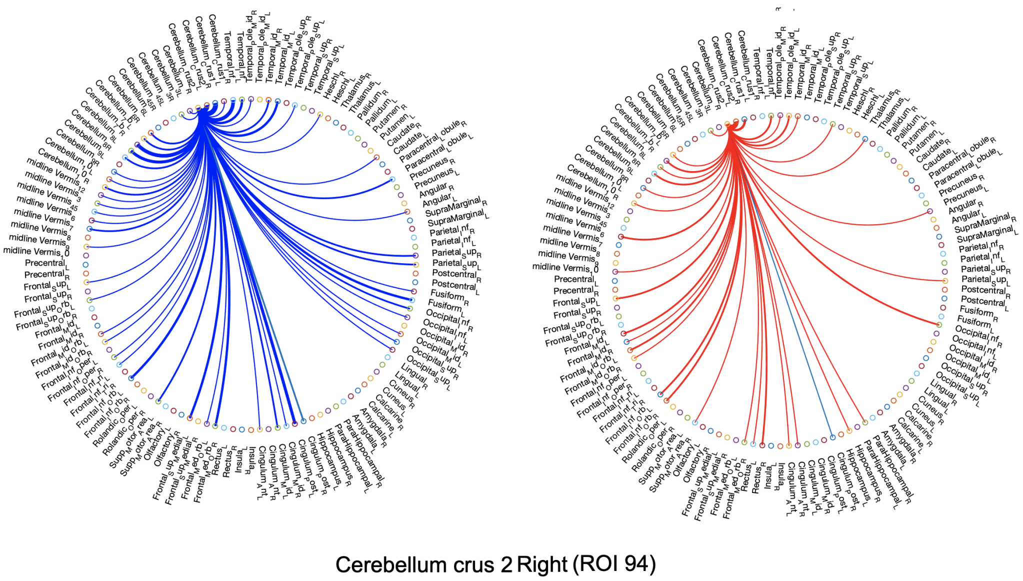


Figure 8: Comparison of average backbone connectivity between right cerebellum crus 2 and other regions between the control and ASD cohort

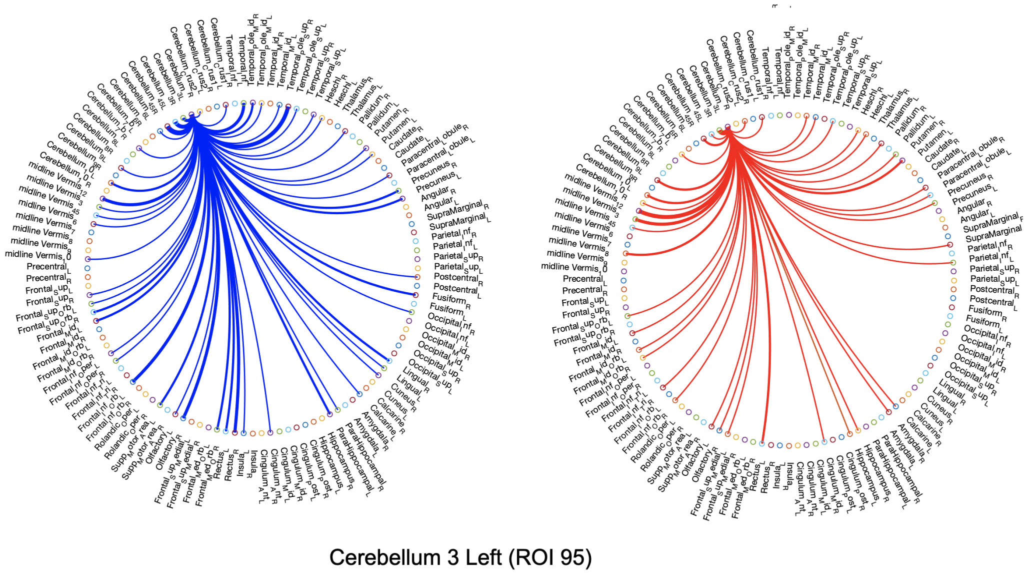


Figure 9: Comparison of average backbone connectivity between left cerebellum area 3 (ROI 95 per AAL) and other regions between the control and ASD cohort

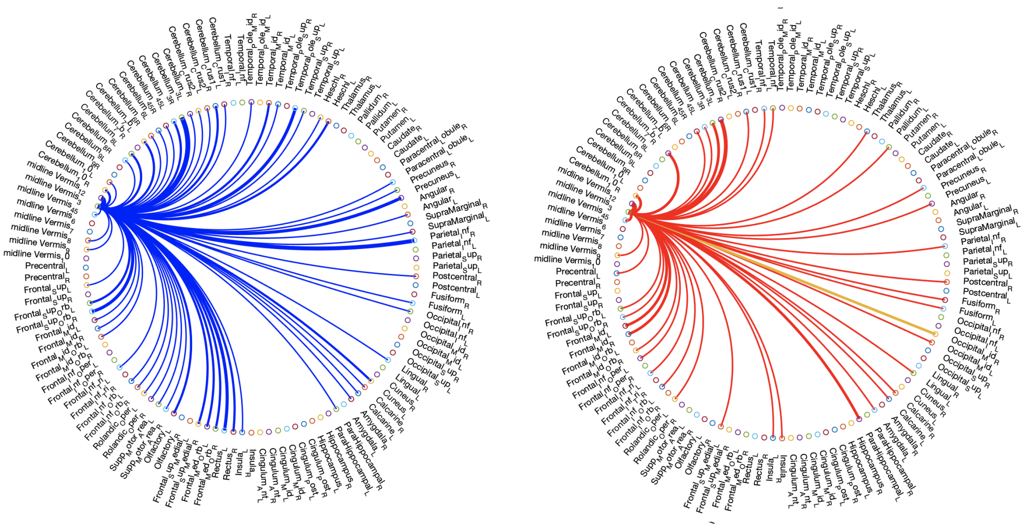

Vermis 3 (ROI 110)

Figure 10: Comparison of average backbone connectivity between the vermis 3 area (ROI 110 per AAL) and other regions between the control and ASD cohort

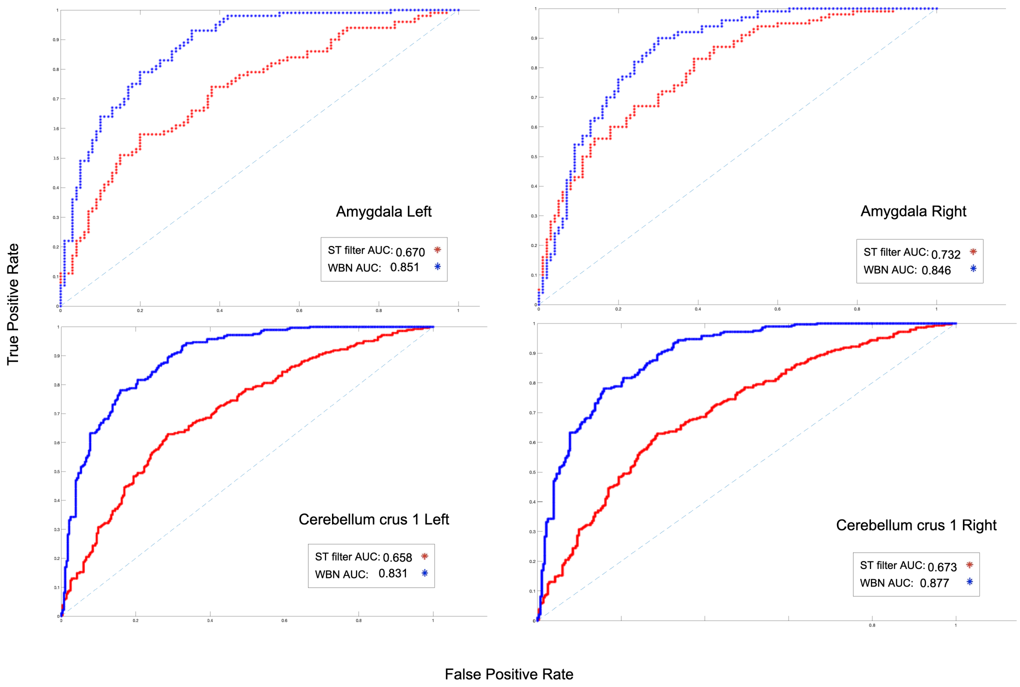


Figure 11: The AUC of detection of injected random weights based on the ST filtering aswell as the proposed approach (WBN) in four different regions where 50 random edges were injected to Amygdalas and 200 random edges were injected to hippocampus areas.
